# Patient-specific "Physical Twin" artery-on-chip platform reveals complex flow-dependent VWF mechanobiology and guides personalized antithrombotic therapy

**DOI:** 10.64898/2026.05.12.724721

**Authors:** Yunduo Charles Zhao, Yanyan Liu, Zihao Wang, Naveen Louis, Nicole Alexis Yap, Arian Nasser, Allan Sun, Yiyao Catherine Chen, Alexander Dupuy, Eugene Marfo Obeng, Mary Meltem Kavurma, Ken S Butcher, Xin Xu, Jingjing You, Freda Passam, Timothy Ang, Lining Arnold Ju

## Abstract

Ischemic stroke triggered by carotid atherosclerosis remains unpredictable because the thrombosis embolization is governed by patient-specific vascular geometry and hemodynamics that cannot be recapitulated in conventional models. Here, we engineered "Physical Twin" artery-on-chip platforms that reproduce individualized carotid anatomy with humanized subendothelial matrix composition and arterial endothelial phenotype under physiological flow patterns. We then established thrombotic microenvironments following laser-induced injury. Computational fluid dynamics-guided experiments across distinct patient geometries reveal that local shear dictates a mechanobiological hierarchy: high-shear bifurcations (∼3,000 s⁻¹) produce von Willebrand factor (VWF) A1-dependent thrombi suppressible by conformationally sensitive inhibitors, whereas low-shear stenoses generate fragile aggregates where VWF-integrin α_IIb_β₃ coupling governs embolization overgrowth. Pulsatile flow suppresses thrombotic growth independent of mean shear. Molecular dynamics simulations reveal why conformationally sensitive nanobodies (caplacizumab) outperform shear-independent aptamers (ARC1172) in high-shear bifurcation flow zones. This Physical Twin platform provides a mechanistic blueprint for geometry-informed, personalized antithrombotic therapy selection.

## 1. Introduction

Ischemic stroke is the second leading cause of death worldwide, with carotid artery atherosclerosis accounting for 15-20% of cases (*1, 2*). Despite advances in imaging and medical management, stroke risk stratification remains imprecise, and nearly 20% of patients on optimal antiplatelet therapy experience recurrent thromboembolic events (*3, 4*). This clinical heterogeneity reflects an incomplete understanding of how patient-specific vascular geometry and local hemodynamics regulate thrombosis initiation, growth, and embolization following plaque rupture or erosion.

Thrombosis at atherosclerotic lesions is fundamentally a mechanobiological process. When endothelial injury exposes the thrombogenic subendothelial matrix, flowing blood subjects adherent platelets to spatially heterogeneous shear forces ranging from <500 s⁻¹ in recirculation zones to >3,000 s⁻¹ at stenotic jets (*5–8*). These mechanical forces drive conformational activation of von Willebrand factor (VWF), an ultra-large multimeric glycoprotein that transitions from a compact, inactive state to an extended conformation capable of capturing platelets via its A1 domain binding to platelet glycoprotein Ibα (GPIbα) (*9–13*). Subsequent thrombus stabilization requires integrin α_IIb_β₃-mediated platelet-platelet cohesion through fibrinogen and VWF C4 domain linkages (*14–16*). However, the relative contributions of these pathways are shaped by local hemodynamics in ways that cannot be predicted from flow magnitude alone.

Existing thrombosis models fail to capture this mechanobiological complexity. Conventional parallel-plate flow chambers impose uniform shear across simplified geometries, neglecting the three-dimensional flow disturbances characteristic of bifurcations and eccentric stenoses (*17, 18*). Animal models, while biologically complete, cannot reproduce human-specific coagulation cascade kinetics or VWF mechanosensing properties (*19, 20*). Recently developed microfluidic systems have begun to overcome these limitations by incorporating physiological shear gradients (*21–23*), yet most rely on generic channel designs that do not reflect patient-specific anatomy, and use venous endothelial cells that are inappropriate for arterial thrombosis (*24–27*), or lack the extracellular matrix (ECM) components essential for VWF-mediated platelet capture under high shear (*28–30*).

The concept of a "Physical Twin"—a microfluidic replica that faithfully reproduces patient-specific anatomy, tissue-level composition, and physiological dynamics—offers a transformative approach to dissecting thrombosis mechanisms and predicting therapeutic responses (*31, 32*). Unlike computational "digital twins" that rely on mathematical models with simplified boundary conditions (*33–35*), Physical Twins preserve geometric complexity while enabling experimental manipulation of molecular pathways in a fully human biological context. For thrombosis, this requires integration of four elements: (i) patient-specific vascular geometry reconstructed from clinical imaging, (ii) arterial endothelial phenotype conditioned under physiological flow, (iii) authentic subendothelial matrix that supports VWF-dependent platelet capture, and (iv) real-time imaging of thrombus dynamics under controlled hemodynamic perturbations.

Recent advances in high-resolution 3D printing now enable fabrication of anatomically accurate microfluidic devices from computed tomography angiography (CTA) data (*30*), while mechanistic insights into VWF conformational regulation have identified discrete molecular epitopes that govern force-sensing (*36–38*). Critically, the VWF A1 domain contains autoinhibitory modules (AIM) comprising N-terminal (N’AIM) and C-terminal (C’AIM) flanking sequences that sterically shield the GPIbα-binding interface in the absence of extensional force (*39–41*). Flow-induced extension peels these regulatory elements, exposing the A1 binding site—a conformational transition that can be selectively interrogated using conformational sensitive antibodies (e.g., 5D2, caplacizumab) versus conformation-insensitive reagents (e.g., ARC1172) (*42, 43*).

Here, we engineer carotid artery Physical Twins that integrate patient-specific geometry, humanized ECM biofunctionalization, human carotid artery endothelial cells (HCtAECs) flow-conditioned to arterial phenotype, and laser-induced injury to establish physiologically authentic thrombotic microenvironments. Combining real-time confocal microscopy, computational fluid dynamics, and molecular dynamics simulations of VWF mechanotransduction, we reveal a shear-dependent mechanobiological hierarchy in which geometry dictates the dominant adhesive pathway, thrombus mechanical stability, and embolic risk. We demonstrate that pulsatile flow waveforms suppress thrombosis formation independent of mean shear, and that rational selection of VWF inhibitors based on conformational gating principles enables geometry-specific therapeutic optimization. This Physical Twin platform provides a mechanistic framework for personalized stroke prevention strategies.

## 2. Results

### 2.1 Engineering the "Physical Twin": Biofabrication and arterial endothelial phenotype under flow conditioning

To establish physiologically authentic thrombotic microenvironments, we developed a fabrication pipeline that preserves patient-specific anatomy while recapitulating arterial endothelial biology and subendothelial matrix composition (Fig. 1). Carotid geometries form 6 patients were reconstructed from clinical CTA scans, segmented using DICOM visualization software, and manually refined to correct for imaging artifacts (Fig. 1A). The clinical characteristics of these patients are summarized in Table 1. Pathological accuracy was confirmed by a Neurologist before exporting stereolithography files. To enable optical imaging, we implemented glass-substrate digital light processing (DLP) 3D printing (*44, 45*) with 10-μm pixel resolution, yielding master molds that preserve bifurcation angles, eccentric stenoses, and local curvature with negligible dimensional error (Fig. S1). Soft lithography with polydimethylsiloxane (PDMS) casting generated elastomeric microchannels mechanically clamped to glass coverslips for confocal-compatible live imaging. For fabrication purposes, the reconstructed vessel geometry was digitally divided into two complementary halves along the medial axial plane. After PDMS replication, these halves were precisely aligned and reassembled to generate a fully enclosed full-lumen microchannel as previously described (*27, 30*).

**Figure 1.**
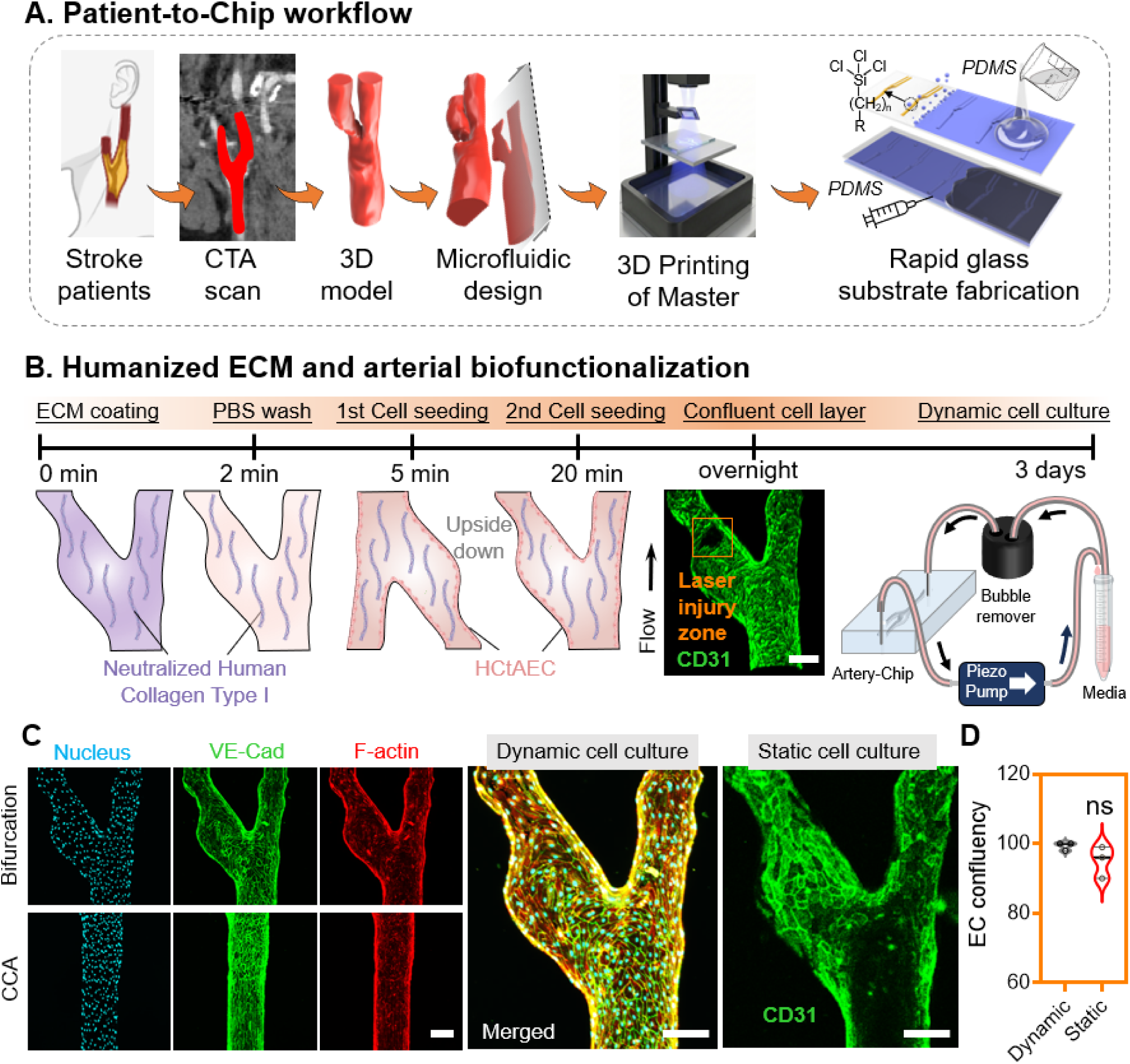
Engineering a patient-specific “Physical Twin” carotid artery–on-chip with humanized matrix and arterial endothelium. **(A)** Patient-to-chip workflow. Clinical CTA data are segmented to reconstruct individualized carotid geometries, which are converted into microfluidic designs and fabricated using glass-substrate digital light processing. PDMS replicas are bonded to functionalized glass to create optically accessible channels that preserve patient-specific bifurcation and stenosis architecture. **(B)** Humanized extracellular matrix and endothelial biofunctionalization. Channels are coated with neutralized human collagen type I, followed by sequential seeding of human carotid artery endothelial cells (HCtAECs) to establish a confluent arterial monolayer. After overnight attachment, constructs undergo dynamic pre-conditioning to promote arterial phenotype prior to thrombosis experiments. Localized laser injury (orange box) is introduced at geometry-informed sites to expose the collagenous subendothelium under controlled shear. Scale bar, 200 μm. **(C)** Arterialization under dynamic culture immunofluorescence demonstrates mature endothelial organization following flow conditioning: continuous VE-cadherin junctions, aligned F-actin stress fibers, and uniform nuclear orientation in both bifurcation and common carotid segments. Compared with static culture, dynamically conditioned cells exhibit elongated morphology and improved cytoskeletal alignment consistent with in vivo arterial endothelium. Scale bar, 200 μm. **(D)** Endothelial confluency assessment. Quantification of CD31-positive coverage shows comparable monolayer confluency between dynamic and static cultures (ns), indicating that flow conditioning enhances structural organization without compromising cell density.

**Table 1.** Summary of Patient Information and Clinical Characteristics. The table consists of key data such as age, sex, and specific details regarding carotid artery pathology. Patient code: Unique identifier for each patient in the study. Age: The patient’s age at the time of diagnosis. Gender: The gender of the patient. Side of Stenosis: Indication of whether the stenosis is present on the left or right side of the carotid artery. Radiology findings: Clinical diagnosis based on CTA and DSA imaging, including the presence of stenosis, plaque characteristics, and ulceration sites. Pre-morbidity medications: List of anticoagulants and antiplatelet agents taken by each patient before the stroke

| Patient code | Age | Sex | Height (cm) | Weight (kg) | Date of Stroke | Side of Stroke* | 3D-recon Segment | DSA * | Symptom | Radiology findings | Pre-Morbidity Medications | Recurrent Stroke |
| --- | --- | --- | --- | --- | --- | --- | --- | --- | --- | --- | --- | --- |
| 1 | 65 | M | 185 | 70 | 08.09.2023 | Left | C1 | Yes | Acute stroke | Fibrocalcification, Ulceration, 60 % stenosis at carotid bifurcation | Antiplatelet: Aspirin | Stroke 5 months earlier at the same territory. XRT† to neck and whole body. |
| 2 | 78 | M | NA | 98 | 20.01.2023 | Right | C1 | Yes | Acute stroke | Fibrocalcification, Ulceration, 40% stenosis | Anticoagulant: Rivaroxaban | No |
| 3 | 67 | M | NA | 101 | 12.12.2022 | Left | C1 | No | Acute stroke | Fibrocalcification, 66% stenosis | Antiplatelet: Aspirin + Clopidogrel (*DAPT†*) | TIA† / stroke two months earlier in the same hemisphere |
| 4 | 71 | M | 174 | 87.7 | 31.08.2022 | Right | C1 | No | Facial asymmetry, dysphasia, dysphagia, dysarthria | < 10% stenosis | None | No |
| 5 | 56 | M | 176 | 83 | 29.03.2023 | Both<br>(*Right as reference*) | C1 | Yes | Acute stroke | Fibrocalcification, Ulceration, 20% stenosis | None | No |
| 6 | 70 | M | NA | NA | 07.09.2022 | Pons | C1 | No | Acute stroke | Fibrocalcification, 10% stenosis | Antiplatelet: Aspirin | No |

\* Side of Stroke: Left, Right, or Both hemispheres. DSA: Digital Subtraction Angiography.
† DAPT: Dual Antiplatelet Therapy. TIA: Transient Ischemic Attack. XRT: Radiation Therapy.

A critical limitation of prior organ-on-chip thrombosis models has been the absence of a physiologically relevant ECM (*28, 29*). Following endothelial denudation in vivo, VWF binds to exposed subendothelial collagen type I and III, forming the molecular substrate for high-shear platelet capture (*46*). While collagen casting and bioprinting can generate bulk 3D constructs, many vascular chips default to substitute other mammalian collagen (e.g. rat tail) to reduce cost, despite known species-dependent differences in platelet and VWF interactions that can confound translational interpretation. In addition, our comparison highlights that commonly used acid-solubilized mammalian collagen can impair endothelial barrier maturation if residual acidity is not completely removed. In our hands, rat-tail collagen coatings (0.1-0.5 mg/mL as suggested by the commercial supplier) supported cell attachment but yielded discontinuous CD31-positive junctions across concentrations, consistent with incomplete formation of a functional monolayer (Fig. S2A). Although neutralized human type I collagen coating (0.1-0.5 mg/mL) better preserved endothelial phenotype, it frequently deposited as heterogeneous, fiber-like structures in confined channels (Fig. S2B), reducing coating uniformity even when junctional organization was maintained.

Here we optimized a flushing-coating strategy that decouples collagen presentation from bulk gelation. Briefly, highly concentrated neutralized human type I collagen (6 mg/mL) is introduced to transiently wet the lumenal surface and is immediately flushed with PBS, producing a thin, stable collagen layer compatible with subsequent endothelialization (Fig. 1B). Importantly, this approach also provides a collagen-rich thrombogenic interface that becomes accessible upon laser injury, mimicking subendothelial matrix exposure during plaque disruption while retaining the reproducibility and optical accessibility of PDMS-based microfluidic systems.

A second unresolved issue in vascular organ-on-chip platforms is the frequent use of human umbilical vein endothelial cells (HUVECs), which exhibit a venous phenotype inappropriate for modeling arterial thrombosis (*24*). Here we used primary HCtAECs seeded onto microchannels and subjected them to 72 hours of arterial-level shear flow (12–15 dyne/cm², corresponding to ∼1,200–1,500 s⁻¹) as a proof-of-concept pre-conditioning regimen (Fig. 1C).

Immunofluorescence imaging before and after flow conditioning revealed a morphological transition (Fig. 1C). In static culture, HCtAECs exhibited a cobblestone morphology with randomly oriented cell-cell junctions. Immunostaining demonstrated continuous VE-cadherin localization at intercellular borders and intracellular F-actin fibers, confirming the establishment of mature adherens junctions and cytoskeletal organization. Flow conditioning further enhanced endothelial confluency (Fig. 1D; Fig. S2C), although this increase did not reach statistical significance.

Following endothelialization, we applied focused laser injury to discrete 50-100 μm diameter regions, creating reproducible endothelial denudation with minimal thermal damage to surrounding cells (Fig. 1B; Fig. S3). Laser ablation locally exposed the underlying collagen matrix. Whole human blood anticoagulated with sodium citrate was recalcified to 2 mM using calcium chloride and labeled with fluorescent anti-CD42b (GPIbα). The blood was then perfused at patient-specific flow rates determined by computational fluid dynamics analysis as previously described(*30*). Real-time confocal microscopy enabled visualization of thrombus dynamics with 6-second temporal resolution over 10-minute observation windows (Videos S1-S3). This integrated Physical Twin platform thus reproduces the key biological and mechanical determinants of post-rupture thrombosis in a fully human, optically accessible format.

### 2.2 Vascular geometry and hemodynamics govern thrombotic phenotype and embolic risk

To investigate how patient-specific anatomy dictates thrombotic behavior, we fabricated Physical Twins from six carotid geometries spanning a spectrum of clinical presentations. Patient 1 (high-grade eccentric stenosis with ipsilateral stroke recurrence), Patient 2 (ulcerated ICA plaque), Patient 3 (moderate stenosis with stroke history), Patient 4 (mild disease), Patient 5 (ulcerated asymptomatic plaque), and Patient 6 (mild disease) (*47, 48*). Computational fluid dynamics (CFD) analysis of each geometry revealed striking heterogeneity in local hemodynamics (Fig. 2A-B; Fig. S4A).

**Figure 2.**
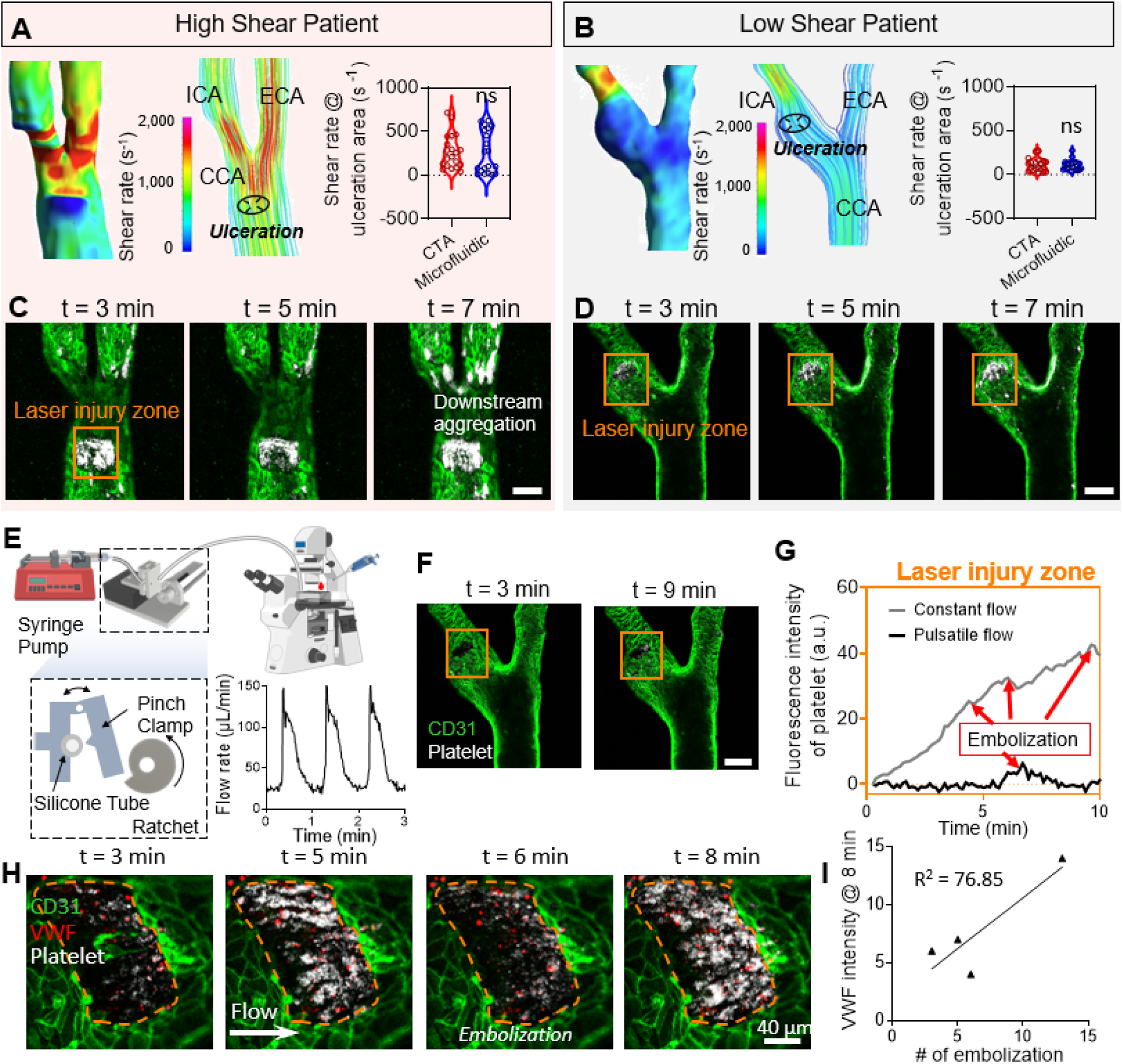
Geometry-driven hemodynamic phenotypes dictate thrombosis versus embolizatio. **(A)** High-shear patient geometry. CFD-derived maps show a focal stenotic jet within the common carotid artery (CCA) producing peak shear rates >3,000 s⁻¹. Streamline analysis along the injury axis reveals a steep proximal–distal shear gradient consistent with force-favored VWF activation. **(B)** Low-shear/disturbed-flow geometry. Shear rate near the ulceration area remains <500 s⁻¹, until reaching to the ICA stenosis, representing a regime permissive to collagen-supported adhesion. **(C–D)** Divergent thrombotic phenotypes. Real-time confocal imaging after laser injury shows that the high-shear patient develops a compact, spatially confined thrombus with progressive downstream aggregation (C), whereas the low-shear patient forms aggregates that remain localized (D). Scale bars, 200 μm. **(E–G)** Effect of pulsatility. A pulsatile perfusion module reproduces physiological velocity waveforms (E). Under pulsatile flow, platelet accumulation at the injury site is markedly reduced and exhibits episodic decreases corresponding to embolization events (F–G), whereas constant flow promotes monotonic growth. Scale bars, 200 μm. **(H)** VWF-rich thrombus architecture. Dual labeling reveals that VWF deposition (red) precedes and colocalizes with platelet aggregates (white) on the endothelial surface (green), with visible shedding of VWF-positive fragments during embolization. Scale bar, 40 μm. **(I)** Embolization correlates with VWF accumulation. Across experiments, VWF fluorescence intensity at 8 min strongly correlates with the number of embolization events (R² = 76.85).

We then selected patients 1 and 2 as the high- and low-shear representatives for mechanobiological investment. Patient 1 exhibited a focal high-shear jet at the bifurcation, with wall shear rates exceeding 3,000 s⁻¹ along streamlines traversing the ulcerated region (Fig. 2A). In contrast, Patient 2 displayed an asymmetric stenotic geometry after the carotid bulb. Specifically, the global flow accelerates as the cross-section narrows downstream of the carotid bulb, creating a locally accelerating flow field, while the shear rate at the endothelial injury site remained within a comparatively low range (∼300–1,000 s⁻¹) relative to the high-shear patient (Fig. 2B). These CFD-derived hemodynamic profiles provided quantitative predictions of the local mechanical forces acting on thrombi, enabling mechanistic interpretation of subsequent experimental observations.

Thrombosis dynamics in the high-shear Patient 1 geometry following laser injury were characterized by rapid, spatially confined platelet accumulation (Fig. 2C, Video S1). Within 3 minutes, a stable aggregate formed at the injury site and exhibited progressive growth with minimal embolization throughout the 10-minute observation period. In contrast, larger thrombi developed at the distal bifurcation approximately 6–7 minutes after perfusion. Embolization events, detected as discrete platelet-rich fragments detaching from the parent thrombus, occurred predominantly at the distal bifurcation rather than at the ulceration site. These observations suggest that the injury site, where both VWF and collagen are exposed, supports mechanically stable thrombus formation at the injury zone even under high shear, whereas regions of intact endothelium lacking exposed collagen are more prone to unstable aggregation and embolization.

In stark contrast, thrombosis in the low-shear, disturbed-flow Patient 2 geometry exhibited more embolization at the injury zone (Fig. 2D, Video S2). Despite achieving comparable peak thrombus size, aggregates underwent frequent embolization, with 3-5 discrete shedding events per 10 minutes. Time-lapse imaging revealed a characteristic "growth-shedding-regrowth" cycle in which platelet accumulation was periodically interrupted by detachment of thrombus fragments. These embolic events produced transient drops in fluorescence intensity, followed by renewed platelet deposition on the residual aggregate. Despite this instability, total platelet accumulation at the laser injury zone over 10 minutes was not significantly different from Patient 1, indicating uncoupling of thrombus size from embolic risk.

To determine whether temporal flow dynamics further modulate thrombus stability, we subjected Patient 2 geometry to 0.5 Hz pulsatile flow versus constant flow at matched mean shear rate (Fig. 2E, Fig. S4B, Video S3). Pulsatile flow dramatically altered thrombotic behavior: thrombi exhibited slower growth rate (Fig. 2F). Quantitative fluorescence analysis revealed a sawtooth pattern, with embolization frequency increasing ∼3-fold under constant flow conditions (Fig. 2G). These data demonstrate that thrombus mechanical stability is sensitive not only to mean shear magnitude but also to temporal shear gradients, with pulsatile suppressing fragmentation independent of average hemodynamic load.

Given the strong association between embolization and VWF accumulation observed in multiple patient geometries, we hypothesized that VWF multimer density at the injury site might predict embolic risk. To test this, we performed dual-labeling experiments with fluorescent anti-VWF and anti-CD42b antibodies, enabling simultaneous visualization of VWF and platelet recruitment (Fig. 2H). VWF accumulation was observed to precede platelet adhesion by ∼60 seconds, consistent with VWF serving as the initial molecular "glue" bridging collagen and platelets (*49, 50*). Linear regression analysis demonstrated a strong positive correlation (R² = 0.77) between VWF fluorescence intensity and embolization frequency (Fig. 2I).

Together, these findings establish that thrombotic phenotypes are emergent properties of vascular geometry and hemodynamics, with high-shear stenoses favoring stable growth at the endothelial injury zone, and low-shear stenoses generating fragile, embolization-prone aggregates. Furthermore, VWF accumulation serves as a biochemical predictor of mechanical instability, providing a molecular readout of embolic risk.

### 2.3 VWF A1 conformational gating governs shear-dependent thrombosis initiation

The observation that thrombotic phenotypes varied dramatically across patient geometries suggested that distinct molecular mechanisms dominate different hemodynamic regimes. VWF is the only adhesive protein uniquely activated by mechanical force: extensional flow unfurls VWF multimers, exposing the A1 domain binding site for platelet GPIbα (*9–11*) (Fig. 3A). However, A1-mediated capture is suppressed in the resting state by an autoinhibitory module (AIM) comprising N-terminal (N’AIM, residues 1238-1271) and C-terminal (C’AIM, residues 1459-1493) flanking sequences that sterically occlude the GPIbα-binding interface (*13, 51, 52*).

**Figure 3.**
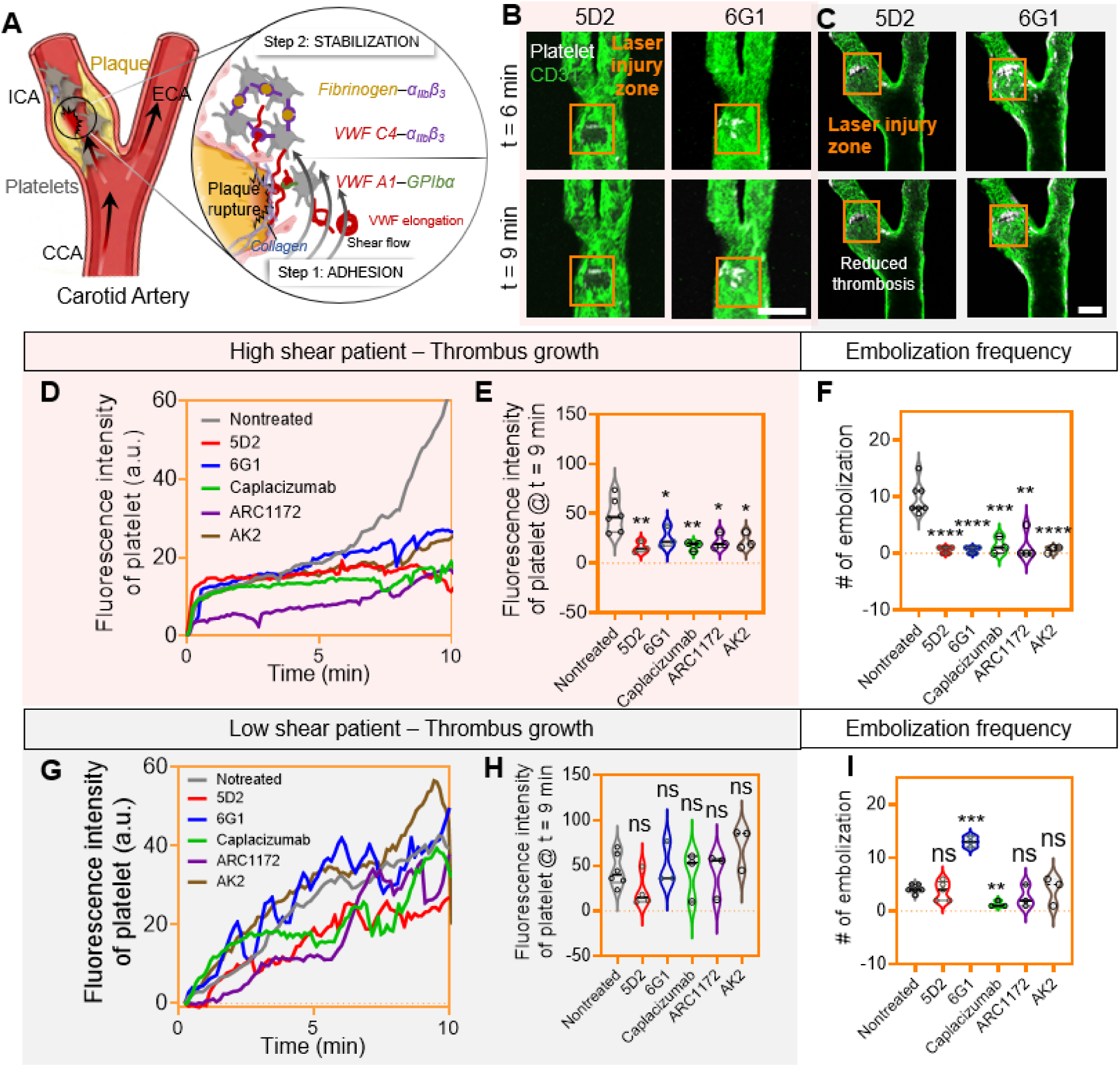
Shear-dependent efficacy of VWF–GPIbα axis inhibitors in high- versus low-shear patients. **(A)** Mechanistic schematic. After laser exposure of collagen at the plaque rupture site, platelet adhesion proceeds via VWF A1–GPIbα capture (Step 1), followed by integrin-mediated stabilization (Step 2) through two competing bridges: fibrinogen–α_IIb_β₃ and VWF-C4–α_IIb_β₃. Inhibitors used: 152B6 (selective disruption of VWF-C4–α_IIb_β₃), LJ-P5 (predominant interference with VWF-α_IIb_β₃ engagement), and tirofiban (blocks fibrinogen binding to α_IIb_β₃). **(B-C)** Representative time-lapse imaging under A1–GPIbα inhibition for the high-shear (B) and low shear (C) geometry. Confocal images at 6 and 9 min after laser injury show platelet (white) accumulation on endothelium (green) within the defined injury zone (orange). Treatments include conformation-sensitive anti-A1 antibody 5D2, conformation-insensitive 6G1, nanobody caplacizumab, aptamer ARC1172, and direct GPIbα blocker AK2. Scale bar, 200 μm. **(D)** Platelet fluorescence intensity over time (high shear). Growth curves confirm that blockade of the A1–GPIbα interaction uniformly attenuates platelet recruitment in the high-shear stenotic environment. **(E)** Endpoint platelet burden at 9 min (high shear). Violin plots show significant reduction for 5D2, 6G1, caplacizumab, ARC1172, and AK2 versus untreated control (*p < 0.05; **p < 0.01), indicating that high-shear thrombosis is critically dependent on force-activated VWF capture. **(F)** Embolization frequency (high shear). All inhibitors significantly decrease embolization relative to control, demonstrating that limiting VWF-mediated tethering improves mechanical stability in stenotic flow (***p < 0.001; ****p < 0.0001). **(G)** Low-shear patient: platelet intensity kinetics. In disturbed/low-shear geometry, inhibition of the A1–GPIbα axis no longer uniformly limits growth. Curves for 6G1, caplacizumab, ARC1172, and AK2 overlap with control, whereas 5D2 shows only partial suppression. **(H)** Endpoint burden in low shear. None of the inhibitors significantly reduce platelet intensity at 9 min (ns), indicating that collagen-supported adhesion and non-VWF pathways sustain thrombus formation when shear forces are modest. **(I)** Embolization in low shear. Drug effects become dissociated from thrombus size. 6G1 markedly increases embolization (***p < 0.001), consistent with AIM destabilization promoting mechanically fragile aggregates, whereas caplacizumab significantly reduces embolization (**p < 0.01). 5D2, ARC1172, and AK2 show no significant change (ns). All data represent independent biological experiments (n ≥ 3) performed using multiple blood donors. Statistical comparisons were performed using a two-tailed Student’s t-test

To dissect whether thrombosis initiation in our platform is gated by VWF–GPIbα engagement, we deployed a mechanistically orthogonal inhibitor panel targeting the VWF A1–GPIbα axis at multiple nodes (Fig. 3B-C; Table 2). 5D2 is a conformational sensitive anti-A1 antibody (*53*), whereas 6G1 binds a linear epitope within the C’AIM region that remains accessible in both compact and extended states (*42*). Caplacizumab (anti-A1 nanobody) and ARC1172 (A1-binding DNA aptamer) inhibit platelet tethering by targeting distinct sites on A1, and AK2 blocks the platelet receptor arm by inhibiting GPIbα. We hypothesized that if high-shear thrombosis is driven by force-unfurled VWF and A1–GPIbα capture, then inhibition at either ligand (A1) or receptor (GPIbα) should suppress both thrombus growth and embolization. Conversely, in low-shear environments where collagen-mediated platelet adhesion can contribute more strongly, blockade of A1–GPIbα may lose efficacy for thrombus growth while retaining selective effects on thrombus stability.

**Table 2:**
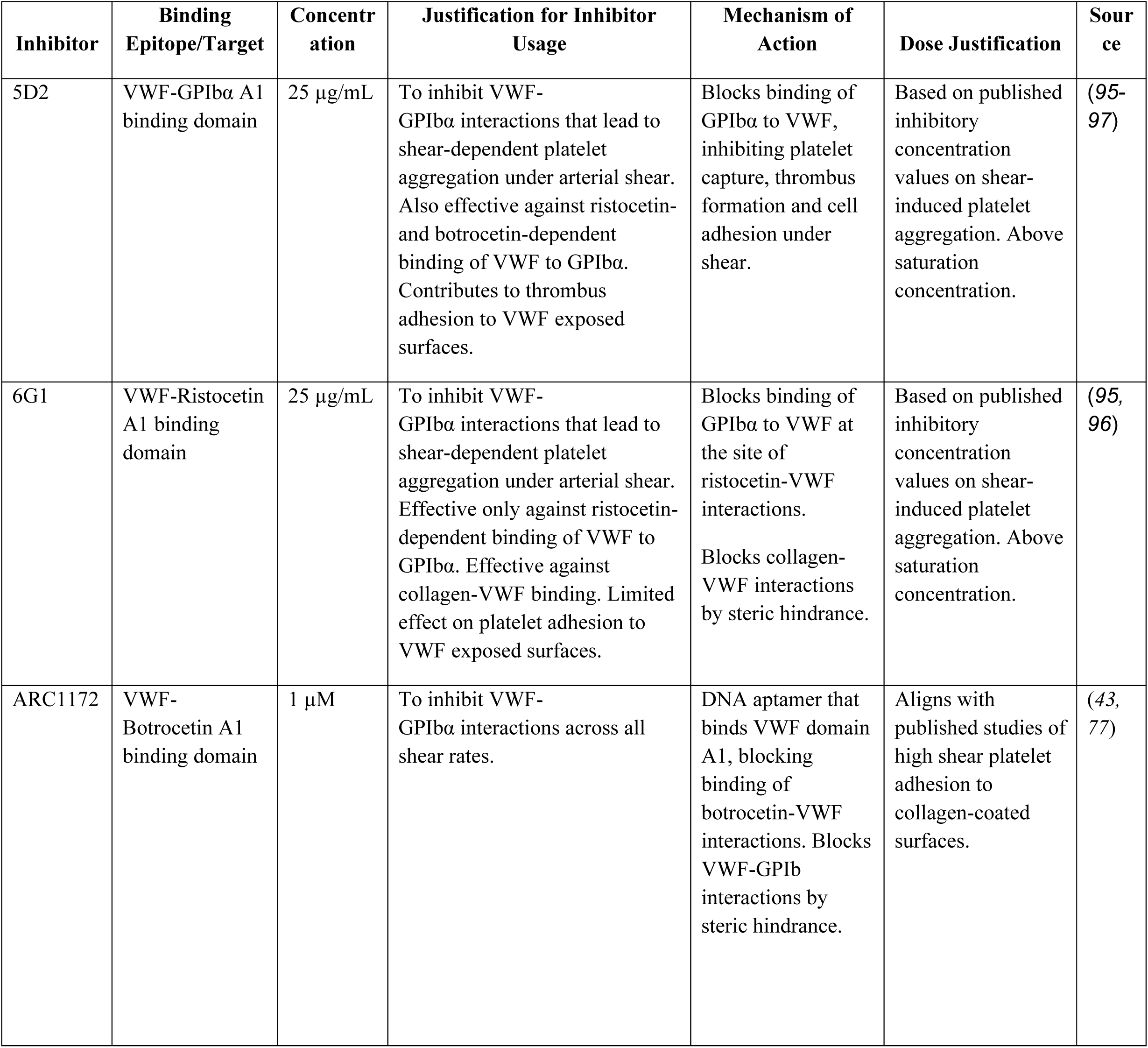
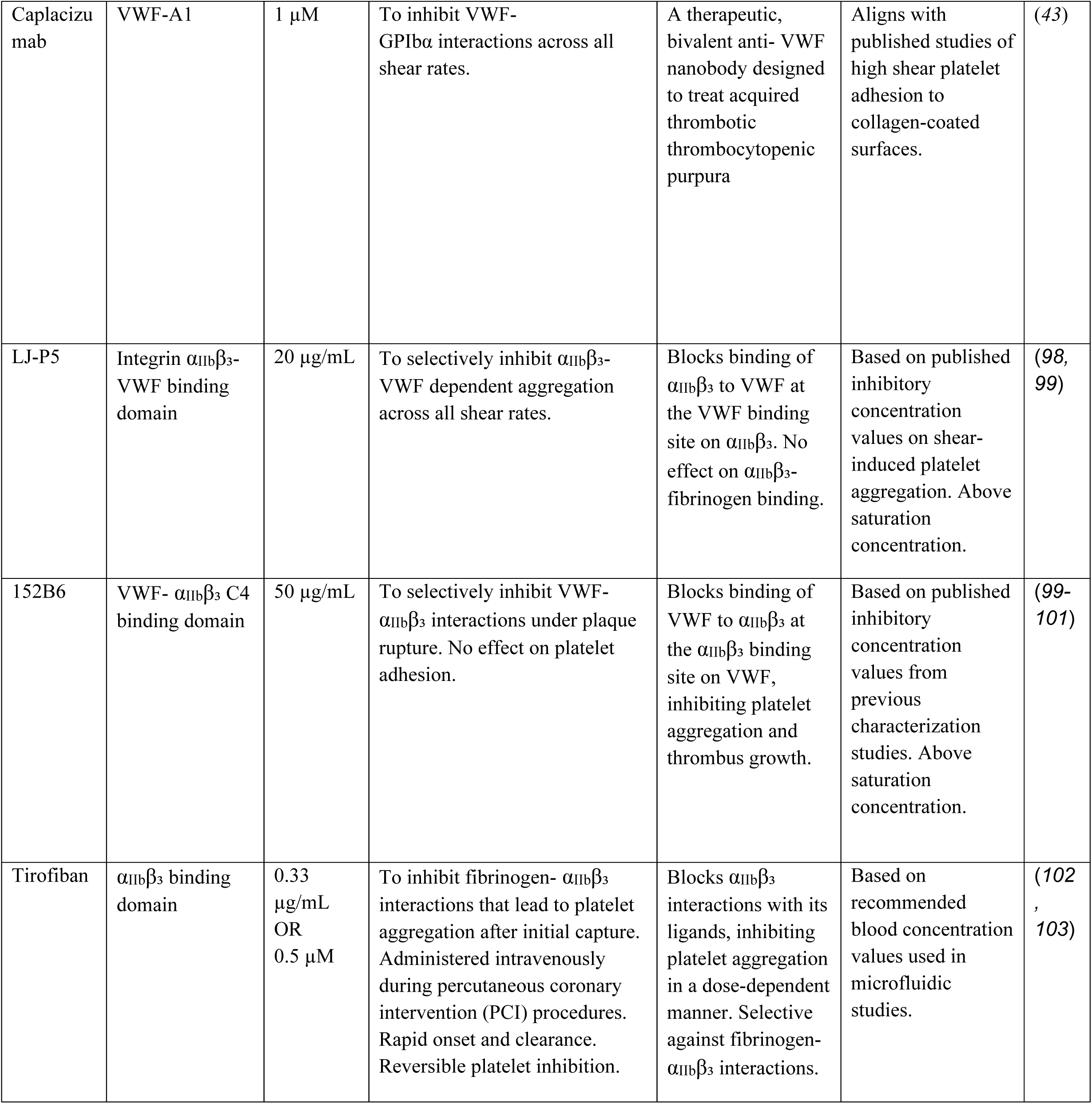
Inhibitor Library & Justification.

In the high-shear patient geometry, time-lapse imaging demonstrated rapid platelet accumulation at the laser injury site in untreated controls (cf. Fig. 2A). Inhibiting the A1–GPIbα pathway at either the VWF side (5D2, 6G1, caplacizumab, ARC1172) or the platelet receptor side (AK2) consistently suppressed thrombus development (Fig. 3B). Quantification of platelet area over time revealed near-complete abrogation of growth across all inhibitor conditions, with residual accumulation only emerging late in the observation window for some treatments (e.g., ARC1172; Fig. S5). Platelet fluorescence intensity traces showed similar pattern, with all inhibitors markedly reducing thrombus accumulation relative to untreated controls (Fig. 3D). Endpoint analysis at 9 min confirmed significant reductions in platelet signal for each intervention (Fig. 3E), establishing that thrombus initiation and expansion under high shear is overwhelmingly dependent on VWF A1–GPIbα capture. Under 6G1 treatment, thrombosis initiation was modestly delayed, and thrombi reached approximately 40% of control size by 9 minutes (Fig. 3E). Although 6G1 reduced thrombus burden relative to untreated conditions, suppression was less pronounced than with 5D2. However, no direct statistical difference was observed between 6G1 and 5D2 in this dataset.

Importantly, embolization frequency was also strongly reduced under high shear (Fig. 3F). All A1–GPIbα perturbations decreased embolization relative to controls, with caplacizumab, 6G1, ARC1172 and AK2 showing particularly low event counts and 5D2 also significantly suppressing embolic shedding. Together, these results define a high-shear regime in which VWF mechanounfolding and A1–GPIbα tethering are the dominant gatekeeper of both thrombus growth and embolization risk.

In the low-shear patient geometry, the same inhibitor panel produced a fundamentally different phenotype. Platelet fluorescence intensity increased over time in all conditions (Fig. 3C and 3G), and endpoint quantification showed no significant reduction in thrombus burden at 9 min for 5D2, 6G1, caplacizumab, ARC1172, or AK2 compared with untreated controls (Fig. 3H). Thus, unlike high shear, thrombus growth in low shear is not obligatorily dependent on the VWF A1–GPIbα axis.

Despite comparable levels of platelet accumulation in the low-shear geometry, the embolization phenotypes diverged markedly among A1–GPIbα pathway inhibitors (Fig. 3I). In this regime, 6G1 treatment significantly increased embolization frequency relative to untreated controls, whereas caplacizumab significantly reduced embolization, and 5D2, ARC1172, and AK2 showed no statistically significant suppression. These results indicate that, under low shear, simply occupying the A1 domain (as 6G1 does) can destabilize nascent platelet aggregates without preventing their formation, leading to increased mechanical fragility and shedding.

Mechanistically, this behavior is consistent with what is known about the VWF AIM and its gating of A1 accessibility. One possible explanation is that 6G1 perturbs the normal autoinhibitory regulation of A1 accessibility, thereby modifying the stability of platelet–VWF contacts under low shear. Published studies have demonstrated that destabilization of the AIM by antibodies or mutations can promote VWF activation via mechanical unfolding of the autoinhibitory module, even under sub-optimal shear (*43*). Under low shear, where collagen-supported adhesion can already sustain platelet recruitment, this partial “AIM loosening” may increase transient, weak adhesive contacts that fragment more easily, thereby increasing embolic shedding even while thrombus size remains unchanged. Thus, while all A1–GPIbα inhibitors converge to suppress high-shear thrombus growth and embolization, their effects under low shear are determined by how each perturbation influences the VWF mechanosensing and downstream aggregate architecture.

### 2.4 Integrin α_IIb_β₃-mediated pathways exhibit shear-dependent hierarchy in thrombus stabilization

While VWF A1 mediates initial platelet capture, sustained thrombus growth and mechanical stability require integrin α_IIb_β₃-dependent platelet-platelet cohesion (*14–16*). Upon platelet activation, αIIbβ₃ serves as a central adhesive hub that can be bridged by two principal ligands: fibrinogen, which binds via its RGD motif; and VWF, which interacts with αIIbβ₃ through its C4 domain (*54, 55*) (cf. Fig. 3A). Although both linkages contribute to aggregate assembly, their relative roles in determining thrombus mechanical integrity under distinct shear regimes have not been systematically defined.

To dissect this hierarchy, we employed three mechanistically distinct α_IIb_β₃-targeting inhibitors: tirofiban, a small-molecule RGD mimetic that selectively blocks fibrinogen binding (*56*); 152B6, a monoclonal VWF antibody that blocks VWF binding to integrin α_IIb_β_3_ but not GPIbα (*57*); and LJ-P5, an antibody that selectively interferes with VWF–α_IIb_β₃ interactions without directly competing with fibrinogen binding, allowing functional separation of VWF-dependent mechanical stabilization from fibrinogen-driven bulk aggregation.(*58*) (Fig. 4A-B, Table 2). We hypothesized that if high shear imposed greater detachment forces, thrombus survival would depend on the stiffest molecular linker (fibrinogen-α_IIb_β₃), whereas under low shear, VWF-α_IIb_β₃ might provide adaptive mechanical reinforcement.

**Figure 4.**
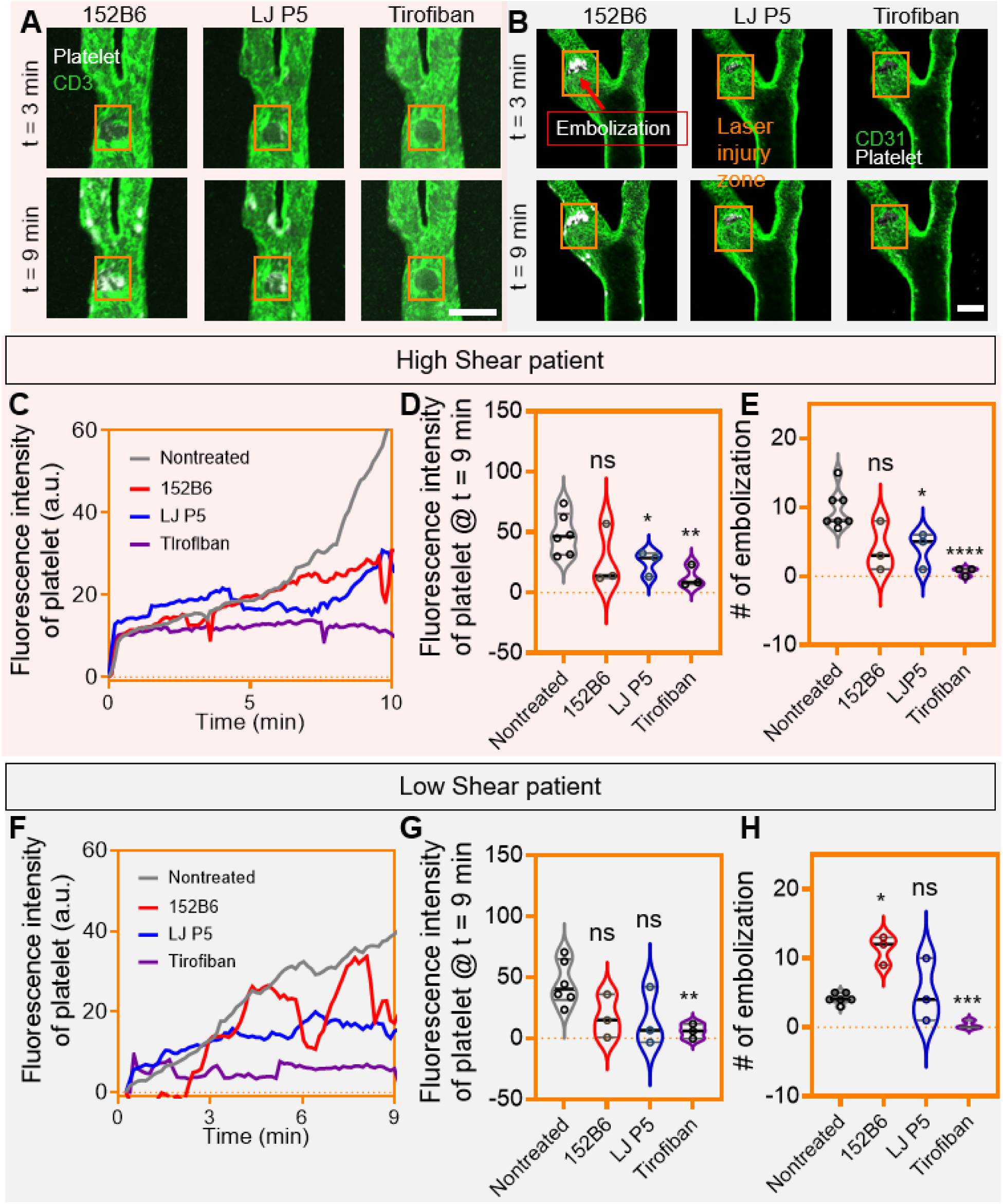
Shear-dependent hierarchy of α_IIb_β₃ ligands: fibrinogen versus VWF in thrombus stability. **(A-B)** Representative imaging for the high (A) and low (B) geometries. Confocal micrographs at 3 and 9 min show platelet aggregates (white) on CD31⁺ endothelium (green) within the injury zone (orange). 152B6 has negligible effect on the high-shear geometry, while produces visible fragmentation/embolization. Tirofiban markedly limits accumulation for both geometries. Scale bar, 200 μm. **High-shear patient (C–E). (C)** Platelet intensity kinetics demonstrate that tirofiban strongly suppresses growth, while 152B6 shows minimal effect and LJ-P5 yields intermediate reduction. **(D)** Endpoint platelet burden at 9 min: tirofiban significantly decreases thrombus size (**p < 0.01), LJ-P5 modestly reduces burden (*p < 0.05), and 152B6 is not significant (ns). **(E)** Embolization counts: tirofiban nearly abolishes embolization (****p < 0.0001); LJ-P5 provides partial protection (*p < 0.05); 152B6 does not differ from control (ns). **Low-shear patient (F–H). (F)** Growth curves show that none of the agents except tirofiban substantially reduce platelet accumulation; 152B6 curves reveal oscillatory drops consistent with repeated fragmentation. **(G)** Endpoint burden: only tirofiban significantly decreases thrombus size (**p < 0.01); 152B6 and LJ-P5 show no reduction (ns). **(H)** Embolization frequency: 152B6 markedly increases embolization (*p < 0.05), producing a “brittle thrombus” phenotype; tirofiban strongly reduces embolization (***p < 0.001); LJ-P5 shows no significant change (ns). We quantitatively define this ’brittle thrombus’ phenotype as an aggregate exhibiting thrombus area comparable to control, combined with significantly increased embolization frequency and platelet intensity fluctuations (see Supplementary Fig. S7).

In the high-shear Patient 1 geometry, tirofiban (0.5 μM) profoundly suppressed thrombosis (Fig. 4A). Thrombus formation was delayed, and aggregates remained small (∼20% of control size) throughout the 10-minute observation period (Fig. 4C-D, Fig. S6). Embolization was nearly abolished, with only rare small fragments detaching (Fig. 4E, p < 0.001). These data indicate that under high shear, fibrinogen-α_IIb_β₃ linkages form the dominant load-bearing scaffold, and their disruption prevents thrombus survival.

LJ-P5 (25 μg/mL) produced significant but more modest reductions in both thrombus burden and embolization (Fig. 4D–E, *p < 0.05). In contrast, 152B6 (25 μg/mL) showed no significant effect on either parameter (Fig. 4D–E, ns). This hierarchy indicates that in high shear, VWF–α_IIb_β₃ interactions are less important, whereas fibrinogen-mediated cohesion is essential for both growth and resistance to embolic failure.

Strikingly different behavior emerged in the low-shear Patient geometry (Fig. 4B). Tirofiban retained robust efficacy, significantly reducing platelet intensity and embolization (Fig. 4F–H, **p < 0.01 and ***p < 0.001), confirming that fibrinogen–α_IIb_β₃ remains the primary driver of bulk aggregation regardless of shear.

However, 152B6 produced a paradoxical phenotype: thrombus size was not reduced (Fig. 4G, ns), yet embolization increased markedly (Fig. 4H, Fig. S7, *p < 0.05, **p < 0.01). Time-lapse imaging revealed repetitive fragmentation of otherwise sizeable aggregates, consistent with mechanically brittle thrombi when the VWF–α_IIb_β₃ stabilizing arm was selectively removed. In contrast, LJ-P5 showed no significant effect on either size or embolization (Fig. 4G–H, ns).

These data reveal a context-specific reorganization of thrombus architecture. Under high shear, fibrinogen–α_IIb_β₃ acts as the dominant load-bearing pathway, rendering VWF–α_IIb_β₃ functionally dispensable. Under low shear, fibrinogen continues to govern bulk growth, but VWF–α_IIb_β₃ becomes a critical mechanical stabilizer, buffering aggregates against flow perturbations. Selective disruption with 152B6 converts thrombi into embolization-prone structures without reducing their mass, uncoupling growth from stability.

This shear-dependent duality explains why inhibitors targeting the same receptor can yield opposite clinical consequences depending on geometry: blockade of fibrinogen binding universally suppresses thrombosis, whereas selective interference with the VWF arm may be protective in high shear yet pro-embolic in low shear. Such findings highlight the necessity of mechanically informed antithrombotic strategies that account for local hemodynamics and extracellular matrix context.

### 2.5 Three-dimensional vascular architecture governs embolization beyond local shear

To test whether local shear magnitude alone was sufficient to explain these phenotypes, we fabricated a simplified single-branch microfluidic model representing only the internal carotid artery of Patient 2, preserving the local shear rate at the injury site while eliminating the bifurcation geometry (Fig. S8A). Despite matched shear at the injury site, thrombus formation in the single-branch geometry exhibited reduced embolization relative to the full bifurcated model (Fig. S8B-C). This demonstrates that three-dimensional vascular architecture, including upstream flow separation and transient recirculation, contributes to mechanical destabilization beyond what can be predicted from local wall shear rate alone. Patient-specific geometry preservation is therefore essential for reproducing clinically relevant embolization phenotypes.

### 2.6 Molecular dynamics simulations reveal mechanisms underlying drug efficacy

To mechanistically explain the divergent efficacies of VWF inhibitors observed in our Physical Twin experiments, we performed flow molecular dynamics (FMD) simulations of the VWF mechanomodule (D′D3–A1–A2–A3) to test whether drug access to VWF is shear dependent. We employed a "water-slice pulling" protocol in which a thin layer of water molecules surrounding the protein is subjected to constant velocity displacement, generating extensional strain analogous to that experienced by VWF in elongational flow (*59*) (Fig. 5A; Video S4). To capture physiologically relevant conformations, both N- and O-linked glycans were incorporated (Fig. S9A), and the system was equilibrated for 500 ns until backbone root mean square deviation (RMSD) convergence was achieved (average RMSD decreased from 0.97 nm at 0–100 ns to 0.38 nm at 400–500 ns; Fig. S9B). The equilibrated structure adopted the compact “bird’s-nest” conformation characteristic of quiescent VWF (*39, 60*)(Fig. S9C).

**Figure 5.**
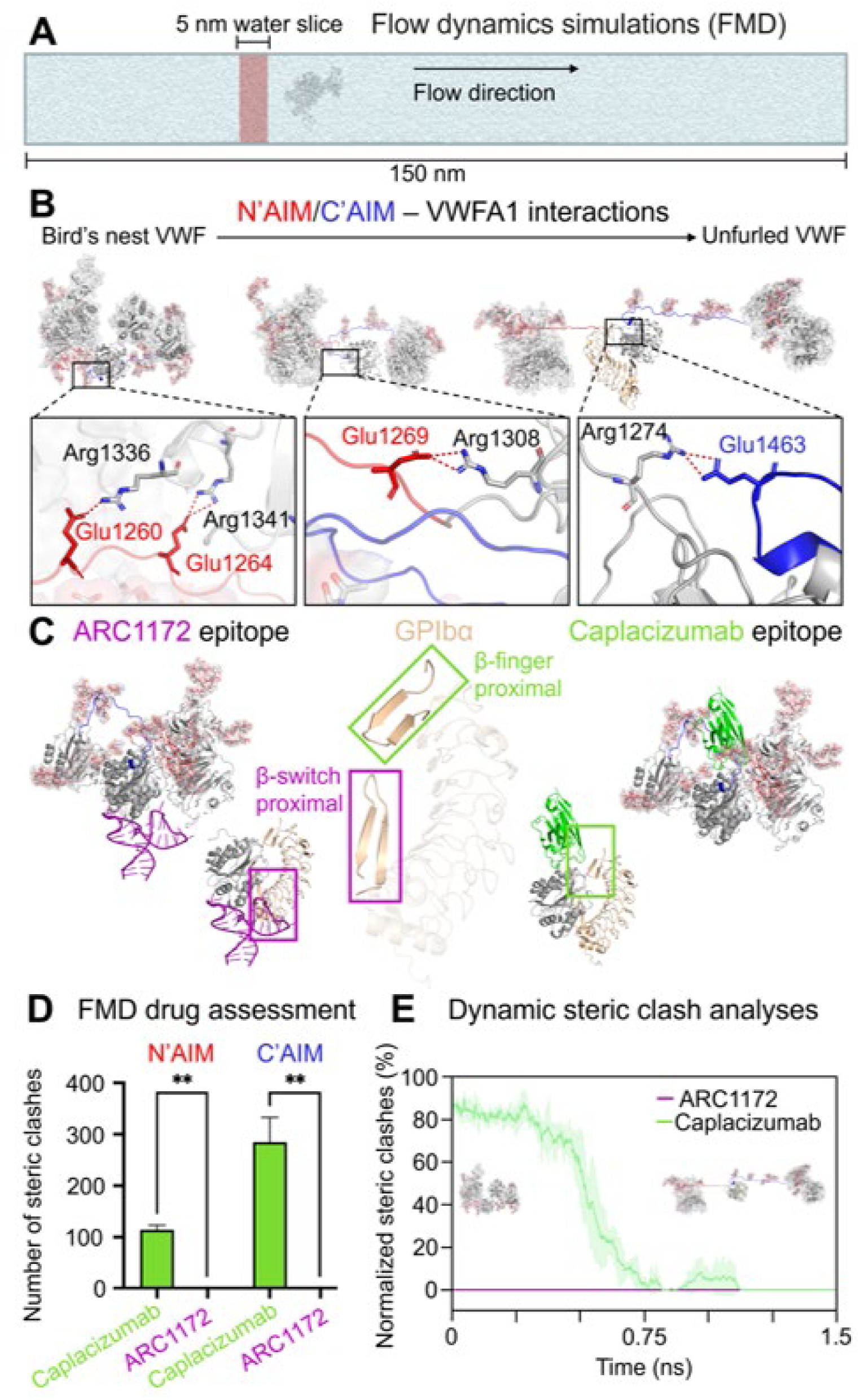
Flow-MD reveals conformational gating of VWF A1 and explains shear-selective drug access. **(A)** Flow dynamics simulation (FMD) framework. Schematic of the water-slice pulling method used to impose extensional flow on the VWF A1 mechanomodule. A 5-nm water slab (red) is translated along the flow direction within a 150-nm periodic box to generate hydrodynamic drag that mimics shear-induced tension. **(B)** Autoinhibitory module (AIM) interactions governing the compact “bird’s nest” state. Top panels: atomic contacts between the N’AIM (red) and C’AIM (blue) segments with the A1 core. Key salt bridges include Glu1260/1264–Arg1336/Arg1341 (left), Glu1269–Arg1308 (middle), and Glu1463–Arg1274 (right), which collectively shield the β-finger region. Bottom panels: structural snapshots along the FMD trajectory showing transition from compact A1 to the unfurled state capable of GPIbα engagement (tan). The boxed region marks the β-finger epitope that becomes solvent-exposed only after AIM displacement. **(C)** Differential accessibility of therapeutic epitopes. Left: ARC1172 (purple) docks adjacent to the β-switch in the compact conformation with minimal steric conflict, indicating that this aptamer can access A1 without prior AIM opening. Right: caplacizumab (green) targets the β-finger–proximal surface that is buried beneath AIM elements in the compact state; docking is favorable only after unfolding. Central inset shows the relative position of GPIbα (tan) for reference. **(D)** Quantified steric clashes from Flow-MD drug assessment. Caplacizumab exhibits a high number of atom-atom clashes with both N’AIM and C’AIM in the compact state, whereas ARC1172 shows minimal conflicts (**p < 0.01), supporting a conformation-independent binding mode for ARC1172 versus a conformation-restricted mode for caplacizumab. **(E)** Dynamic steric-clash trajectories. Normalized clash percentage over time demonstrates that caplacizumab conflicts resolve only after AIM displacement (∼0.6–0.8 ns), while ARC1172 remains stably accommodated throughout the simulation.

In the absence of force, the A1 domain adopted a compact "bird’s nest" conformation, in which N’AIM and C’AIM fold across the β-finger–GPIbα-binding surface, creating a steric barrier to platelet engagement (Fig. 5B, bottom left). Flow triggered progressive loss of interdomain(*61, 62*) and intermodular interactions (Video S4A)(*39*), facilitated by the disruption of salt bridges between the N- and C-terminal autoinhibitory modules (N’AIM and C’AIM), leading to stepwise uncoiling of the mechanomodule (Fig. 5B, top). Under simulated flow extension, these AIM elements disengaged, producing an unfurled A1 configuration compatible with GPIbα engagement (Fig. 5B, bottom right).

To predict drug-binding modes, we performed molecular docking of ARC1172 and caplacizumab to both compact and extended A1 conformations. Mapping the binding epitopes revealed that ARC1172 engages the A1 domain proximal to the GPIbα β-switch, whereas caplacizumab binds near the β-finger region (*63, 64*)(Fig. 5C).

Steric clash analysis showed that ARC1172 remained accessible towards the VWF A1 domain across all conformational states, whereas caplacizumab exhibited steric occlusion by N’AIM and C’AIM in the compact conformation (Fig. 5D). Upon flow-induced unfolding, transient exposure of the VWFA1 domain eliminated these clashes, enabling caplacizumab access to its epitope (Fig. 5E, Video S4B).

These simulations explain why caplacizumab shows high efficacy in high-shear stenotic regions (where VWF is constitutively extended) but reduced potency in low-shear, disturbed-flow zones (where VWF remains partially compact) (*65*). Thus, the AIM acts as a mechanical gatekeeper that differentially licenses inhibitor access to A1, linking molecular structure to the geometry-dependent drug responses observed in the Physical Twin platform.

### 2.7 Personalized mechanomedicine: Geometry-guided therapeutic stratification

Having established that thrombotic phenotype and drug efficacy are dictated by patient-specific hemodynamics, we sought to derive a framework for geometry-guided therapeutic stratification (Fig. 6). We compiled thrombosis and embolization data under treatment with eight mechanistically distinct inhibitors (Fig. 6A).

**Figure 6.**
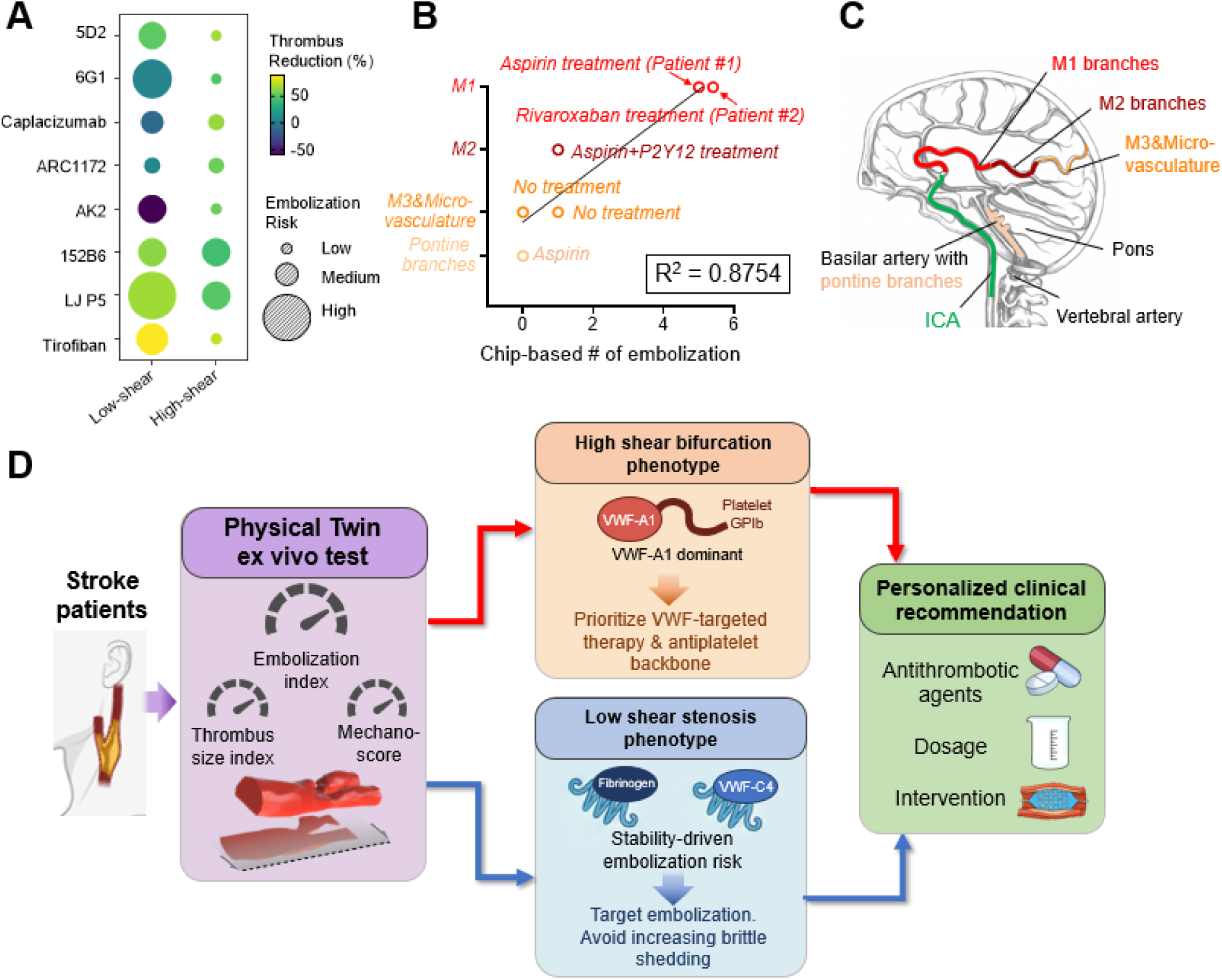
Geometry-guided mechanophenotyping enables a patient-specific antithrombotic decision framework. **(A)** Cross-geometry drug response matrix. Bubble plot summarizing the effect of eight mechanistically distinct inhibitors on thrombus burden (color scale: % reduction) and embolization risk (bubble size) in low-shear versus high-shear phenotypes. The map highlights three functional drug classes: (i) VWF-A1/GPIbα-directed agents (5D2, 6G1, caplacizumab, ARC1172, AK2), (ii) VWF–α_IIb_β₃ modulators (152B6, LJ-P5), and (iii) fibrinogen–α_IIb_β₃ blockade (tirofiban). Responses diverge sharply between shear regimes, revealing that the same drug can be protective in one geometry yet neutral or destabilizing in another. **(B)** Chip embolization predicts clinical severity. Correlation between the chip-derived number of embolization events and the anatomical territory of prior stroke in individual patients. Lesions involving the M1 segment (largest cortical territory; highest clinical impact) correspond to the highest embolization counts, whereas M2 and M3/microvascular territories show progressively lower values. The strong association (R² = 0.8754) supports embolization frequency as a functional surrogate of real-world stroke severity. Treatment histories (aspirin, P2Y12, rivaroxaban, or no therapy) are annotated. **(C)** Vascular territories at risk. Schematic mapping of emboli-sensitive regions in the anterior and posterior circulation, illustrating why identical plaque morphologies can yield disparate neurological outcomes depending on downstream target territory (M1 > M2 > M3/microvasculature; pontine branches). **(D)** Proposed Physical-Twin–guided clinical algorithm. Left: CTA-derived Physical Twin classifies patients into high-shear stenosis or low-shear disturbed-flow mechanophenotypes. Middle: Each phenotype is linked to dominant molecular drivers identified in this study—VWF-dependent capture and fibrinogen-α_IIb_β₃ growth in high shear versus stability-driven embolization risk in low shear. Right: Outputs inform personalized recommendations for agent selection, dosing intensity, and the need for interventional procedures.

Hierarchical clustering of drug responses revealed three patient classes: (1) High-Shear Responders (Patient 1 and 3): strong suppression by VWF A1 inhibitors and tirofiban; minimal effect of 152B6. (2) Disturbed-Flow/Embolic (Patient 2 and 5): strong suppression by ARC1172 and 152B6 in embolization (but not size); tirofiban effective for both. (3) Low-Risk/Stable (Patient 4 and 6): modest thrombus burden; all inhibitors effective, suggesting redundancy in adhesive pathways when hemodynamic stress is low.

To test whether the chip-derived embolization metric reflects clinically meaningful severity, we compared the chip-based number of embolization events with the clinical vascular territory affected in each patient (Fig. 6B). Patients with more severe, proximal large-territory involvement—particularly those with M1 occlusion—exhibited the highest embolization counts on-chip, whereas patients with embolic signatures confined to more distal branches (M2) or limited microvascular territories exhibited lower values. Across the cohort, chip-based embolization frequency strongly correlated with clinical severity stratified by vascular territory (R² = 0.8754, Fig. 6B), supporting the concept that the Physical Twin platform captures a patient-specific propensity for producing emboli capable of causing high-impact arterial occlusions.

To anatomically contextualize this relationship, we mapped the major downstream territories and their relative clinical consequences onto a schematic of the intracranial circulation (Fig. 6C). Emboli entering the M1 segment threaten the largest cortical perfusion territory, producing the greatest ischemic burden, whereas emboli lodging in M2 branches affect a smaller downstream region and emboli reaching M3/microvasculature generally have the most localized territorial impact. The alignment between chip-measured embolization frequency and these anatomical risk gradients indicates that the on-chip embolization phenotype demonstrate a clinically interpretable indicator of emboli generation with potential to cause large-vessel, high-burden stroke.

Integrating the geometry-dependent mechanisms defined above, we formulated a proof-of-concept Physical-Twin–guided decision framework that converts CTA anatomy into functional treatment guidance (Fig. 6D). Patient-specific flow signatures first classify arteries into high-shear stenosis or low-shear disturbed-flow mechanophenotypes. High-shear lesions are characterized by VWF-dependent platelet capture and fibrinogen–α_IIb_β₃–driven growth. In this setting, VWF-A1/GPIbα blockade and fibrinogen-targeted antiplatelet therapy most effectively suppress both thrombus expansion and embolization. In contrast, low-shear geometries exhibit collagen-permissive adhesion and a distinct vulnerability to stability failure, where selective disruption of VWF–α_IIb_β₃ coupling can paradoxically increase embolic shedding despite unchanged thrombus size. The ex vivo Physical Twin assay quantifies these behaviors through an embolization index, thrombus-size index, and composite mechano-score, enabling head-to-head comparison of candidate regimens before clinical selection. Of note, the observed correlation between chip-derived embolization and clinical severity may be sensitive to individual data points and should be interpreted cautiously, e.g. extended blood perfusion (Fig. S10). This workflow holds the potential to establish a functional bridge between imaging and therapy, representing a paradigm shift from one-size-fits-all antiplatelet therapy to geometry-encoded, mechanistically stratified stroke prevention. When anatomy alone is equivocal, chip-based testing provides patient-specific evidence to guide drug choice, dosing intensity, or the need for endovascular intervention (Fig. 6D).

## 3. Discussion

We engineered carotid artery “Physical Twins” that integrate patient-specific geometry, collagenous subendothelium, arterial endothelium, and physiological flow in a humanized system. The incorporation of collagen-matrix biofunctionalization and flow-conditioned HCtAECs directly addresses prior critiques of organ-on-chip thrombosis models (*28, 29*). Collagen type I is essential for VWF immobilization and force-dependent unfolding under arterial shear (*46*); its absence in earlier platforms likely explains why some studies reported VWF-independent thrombosis (*66*). Similarly, the use of HUVECs in arterial flow models has been questioned (*24*), as arterial and venous endothelia differ in mechanosensor expression, VWF secretion kinetics, and inflammatory responses (*67*).

Our Physical Twin platform addresses critical limitations of existing thrombosis models. Unlike parallel-plate chambers that impose uniform shear (*17, 18*), or generic stenosis models with idealized geometries (*21–23*), our approach preserves three-dimensional flow disturbances including bifurcation, stenosis, jet impingement, and transient flow reversal, which emerge only from patient-specific anatomy. The demonstration that a single-branch ICA model fails to reproduce embolization phenotypes observed in the full bifurcated geometry (Fig. S8) underscores that local shear magnitude alone is insufficient; spatial context matters. This finding has important implications for hemodynamic-based risk prediction, suggesting that current clinical algorithms relying on stenosis degree or wall shear stress alone (*68, 69*) may miss geometry-encoded embolic risk.

In addition, we found that vascular architecture establishes a shear-dependent division of labor within platelet adhesion. In high-shear bifurcation, thrombi are stabilized predominantly by fibrinogen–α_IIb_β₃ linkages, while in low-shear stenoses, VWF–α_IIb_β₃ interactions become critical for mechanical stability rather than mass. The paradoxical increase in embolization with 152B6 under low shear illustrates that selective disruption of the VWF–α_IIb_β₃ arm can convert otherwise large aggregates into brittle, embolization-prone structures without reducing size.

The integration of molecular dynamics simulations with microfluidic validation represents a bidirectional "theory-experiment" loop that is rare in organ-on-chip research. FMD revealed that N’AIM/C’AIM shielding restricts access to the β-finger region in the compact state, whereas the ARC1172 epitope remains partially accessible. The data caution that inhibitors targeting similar domains may have opposite embolic consequences depending on how they perturb the AIM–A1 equilibrium, as exemplified by the destabilizing effect of 6G1 in low shear. This structure-function mapping enables rational inhibitor design. For example, next-generation VWF inhibitors could target the β-switch region (like ARC1172) while incorporating RGD-mimetic moieties to simultaneously disrupt α_IIb_β₃ binding.

Our study has several limitations. First, our laser injury model creates instantaneous, full-thickness endothelial denudation, whereas plaque erosion in vivo may involve partial denudation with preserved glycocalyx or apoptotic endothelial remnants (*70*). Incorporating graded injury models would better capture inflammatory priming. Second, our patient cohort (n=6) spans a range of geometries but lacks statistical power for clinical outcome correlation. Third, while our CFD simulations capture gross hemodynamics, near-wall red blood cell dynamics (cell-free layer thickness, hematocrit gradients) influence platelet margination and activation (*71, 72*). Future platforms incorporating physiological hematocrit (∼40%) in bifurcation geometries may reveal hematocrit-shear coupling effects. Finally, our study focused on VWF and α_IIb_β₃ pathways; other adhesive mechanisms (collagen-GPVI, P-selectin-PSGL-1) likely contribute, particularly under inflammatory conditions (*73, 74*).

Clinically, our findings support a paradigm shift toward geometry-guided antithrombotic therapy. Current guidelines stratify stroke risk by stenosis severity (e.g., NASCET criteria) (*75*), yet our data show that two patients with equivalent stenosis can exhibit opposite embolic phenotypes depending on their complex three-dimensional vasculature shape. With caplacizumab recently on market for thrombotic thrombocytopenic purpura (*76*) and ARC1172 in clinical trials (*77*), Physical Twin platforms could enable prospective selection of optimal VWF inhibitor based on preoperative CTA imaging—a true "precision thrombosis medicine" approach.

## 4. Methods

### 4.1 Patient imaging and geometry reconstruction

Six patients with documented cases of carotid artery-related stroke at the Royal Princes Alfred Hospital (RPAH) were selected and deidentified for this study (see Table 1). The inclusion criteria were based on the availability of high-quality CTA and DSA imaging data (New South Wales Telestroke Service) (*78*), essential for accurate 3D reconstruction and hemodynamic analysis. Patients with severe systemic diseases, coagulation disorders, or previous carotid artery interventions were excluded to ensure a homogeneous study population.

Clinical data were collected in accordance with ethical guidelines and patient consent protocols approved by the Sydney Local Health District Human Research Ethics Committee (HREC) – RPAH Zone (X23-0267 & 2023/ETH01607). The collected data included demographic information (age, gender, and relevant medical history) as well as high-resolution CTA and DSA images for each patient. Each patient’s thrombotic events were classified into either comprehensive global thrombosis or localized thrombosis based on both imaging and clinical findings.

De-identified carotid CTA scans (n=6 patients) were obtained under approved protocols with informed consent. DICOM files were imported into SimVascular-23.03.27 (*79*) for segmentation using semi-automated region-growing algorithms. Manual refinement in ANSYS SpaceClaim (v2023) corrected for partial volume artifacts and ensured smooth vessel walls as previously described (*30*). A board-certified neurologist validated anatomical accuracy and pathological features (stenosis eccentricity, ulceration presence). Final geometries were exported as STL files.

### 4.2 Computational fluid dynamics

CFD simulations were performed in ANSYS Fluent (v2023 R2) assuming incompressible Newtonian flow (μ = 3.5 cP, ρ = 1060 kg/m³), as Reynolds numbers derived from measured inlet velocities and vessel diameters for both in vitro and in vivo conditions, yielding values of Re ≈ 3.0 and Re ≈ 670, respectively, which are well below the conventional turbulent threshold of ∼2000. Inlet boundary conditions applied using published velocity profiles derived from clinical data representing the average in vivo wall shear rates produced in the carotid artery measured clinically(*80, 81*) . Pulsatile time-dependent simulation inlet velocities were set based on the measured flow velocities of the syringe pump (Fig. 1E), as a transient table modelling 10 full cycles. Outlet was set to zero pressure. No-slip conditions were applied at walls. The mesh consisted of 1-3 million tetrahedral elements (boundary layer refinement: 5 layers). Simulations solved Navier-Stokes equations utilizing SIMPLE algorithm. Convergence was considered as 10^−6^ for the continuity and momentum equations. Wall shear rate (γ = τ/μ) and velocity were extracted across the entire geometry and plotwith CFD-post. Based on the instantaneous WSS vectors, the time-averaged wall shear stress (TAWSS) and mean oscillatory shear index (OSI) through cycles were computed as

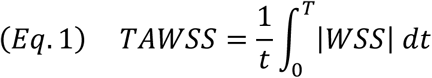

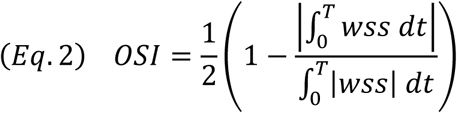

### 4.3 Fabrication of Physical Twin vessel-on-chip devices

The fabrication of the patient-specific carotid artery-on-chip device was conducted according to previously established protocols (*30*).

#### Digital design

The reconstructed blood vessel was bisected along the medial axial plane (Figure 1A). The resulting geometry was arrayed (n=5) with 15 mm spacing along the longitudinal axis of the design space. To facilitate fluidic interfacing, the Common, Internal, and External Carotid Arteries (CCA, ICA, ECA) were extended with cylindrical conduits, the dimensions of which were optimized via CFD simulations to preserve physiological flow characteristics. To ensure precise mold leveling during fabrication, four hemispherical spacers (230 µm height) were integrated into the design substrate. This hemispherical geometry was selected to mitigate detachment risks. The finalized designs were sliced at a 5 µm layer thickness using Voxeldance Additive software and exported as 8-bit PNG sequences. The geometric downscaling factor from CTA to microfluidic scale was approximately 1:20 to 1:30. Following fabrication, the two complementary PDMS halves were aligned and permanently bonded to reconstruct a fully enclosed, full-lumen microchannel. All computational fluid dynamics (CFD) simulations were performed on the complete full-lumen vessel geometry.

#### Glass-Based Printing Substrate Surface Treatment

Glass slides served as the printing substrate and were chemically treated to enhance resin adhesion. The cleaning protocol involved a 5-minute sonication in isopropanol, followed by compressed air drying and a subsequent 5-minute sonication in Milli-Q water. Resin adhesion reinforcement was achieved by immersing the slides for 2 hours in a silanization solution comprising anhydrous Toluene (99.8%) and 3-(Trimethoxysilyl) propyl methacrylate (TMSPM, 98%) at a 90:10 w/w ratio. After rinsed with ethanol and dried under a nitrogen stream, a thin layer of resin (BMF, HT resin) was applied on one side of the glass slide, followed by UV curing. At this stage, the glass printing substrate is ready for high resolution 3D printing. All reagents were sourced from Sigma-Aldrich.

#### High-Resolution 3D Printing

Fabrication was performed using a BMF S240 DLP 3D printer (projection micro-stereolithography). To align the print with the substrate, the STL design incorporated a custom alignment frame consisting of two L-shaped brackets. The glass slide was immobilized on the build platform by UV-curing resin droplets at the bracket interface. Printing utilized a stitch mode to accommodate a 100 × 100 mm² fabrication area at 10 µm pixel resolution, with the print field manually offset to prevent stitch lines from intersecting the vascular features. A variable exposure strategy was employed to optimize both adhesion and feature resolution: Base Layer (Layer 1): High energy exposure (1.5 s at 60 mW/cm²) to secure the construct to the glass. Transition Layers (Layers 2–5): Intensity reduced to 45 mW/cm² to balance adhesion with resolution. Main Structure: Exposure time reduced to 1.0 s (at 45 mW/cm²) to maximize throughput and detail. Surface Features (Final 5 layers): Exposure time increased to 1.5 s to resolve fine geometrical details. Layer height was maintained at 5 µm through a mechanical cycle involving a 4 mm stage retraction and a 3.995 mm return, incorporating a 300-second dwell time between layers to ensure resin relaxation and thickness uniformity.

#### Soft Lithography and Device Assembly

Post-fabrication, the printed molds were detached, washed in 100% ethanol, and subjected to a secondary cure at 50°C for 60 minutes (Form Cure, Formlabs). To facilitate PDMS release, mold surfaces were passivated via vacuum vapor deposition of Trichloro(1H,1H,2H,2H-perfluorooctyl) silane (Sigma-Aldrich).

The microfluidic channels were cast using Polydimethylsiloxane (PDMS, Sylgard 184, Dow Corning) mixed at a 10:1 base-to-agent ratio. An aluminum frame was utilized to control the chip thickness during casting. The polymer was degassed and thermally cured at 70°C for 2 hours. Access ports were created using biopsy punches (6 mm for inlets, 1 mm for outlets).

Final assembly was secured using a custom-printed mechanical clamping system comprising a Rigid 4k resin top plate and a Rigid 10k resin base plate (Formlabs). Prior to experimental use, the assembled devices were sterilized with 80% ethanol and dried using nitrogen gas.

### 4.4 Endothelial cell culture and flow conditioning

Prior to collagen coating, microfluidic chips were sterilized by exposure to ultraviolet (UV) light for 30 min in a biosafety cabinet. For channel coating, 5 μL of type I collagen derived from human sources (6 mg/mL stock solution) was injected into each channel to ensure full coverage of the channel surface. The chips were then gently rinsed with PBS to remove excess collagen prior to cell seeding.

Human carotid artery endothelial cells (HCtAECs; Sigma-Aldrich, 301405A) were cultured in endothelial growth medium (EGM™-2 BulletKit™, Lonza, CC-3162) and maintained at 37 °C in 5% CO₂. Cells between passages 3 and 8 were used for all experiments. Upon reaching 80–90% confluency, cells were detached using TrypLE™ Express enzyme (Thermo Fisher Scientific, 12604013), and resuspended in fresh EGM-2 medium at approximately 1 × 10⁷ cells/mL. For cells seeding, approximately 8 μL of the cell suspension was injected into each channel through the outlet, and excess cell suspension at the inlet was carefully aspirated and collected. The chip was then incubated at 37 °C in 5% CO₂ for 20 min to allow initial cell attachment. Subsequently, an additional 8 μL of cell suspension was injected, and the microfluidic chip was immediately inverted and incubated under the same conditions for 20 min to facilitate cell attachment to the upper surface of the channel. Following cell attachment, 20 μL of EGM-2 medium was gently injected through the outlet to remove non-adherent cells. Fresh medium was then added to the inlet reservoirs, and the chip was cultured overnight (typically 16–18 h) at 37 °C in 5% CO₂, resulting in endothelialization of the channel.

#### Flow pre-conditioning

Culture medium was circulated through the chips using a Bartels mp6 piezoelectric diaphragm micropump (Bartels Mikrotechnik), integrated into a closed-loop perfusion system at 5-10 dyne/cm² (calculated from CFD for a representative ICA cross-section) for 72 h. Pulsatile flow was generated using the same pump operating at 1 Hz, with the flow profile measured via a Fluigent FS6D inline sensor. At the microfluidic scale, the Reynolds number (Re ≈ 3.0) confirmed fully laminar flow, and the Womersley number (Wo ≈ 0.24) indicated the pulsatile flow operated in the quasi-steady regime.

#### Immunofluorescence

After flow, channels were fixed and stained for VE-cadherin (Cell Signaling 2500, 1:100), ZO-1 (Invitrogen 61-7300, 1:100), eNOS (BD 610297, 1:200), KLF2 (R&D Systems AF5889, 1:100), and VWF (Dako A0082, 1:400). Nuclei: DAPI. Imaging: Zeiss LSM 880 confocal. Quantification: ImageJ (alignment angle distribution, junction continuity index).

### 4.5 Laser-induced injury and thrombosis imaging

Focal endothelial disruption was introduced using a confocal-coupled laser ablation system to initiate site-specific thrombosis within the Carotid Artery-Chip. A UGA-42 Caliburn module (355-nm pulsed DPSS; 42 µJ per pulse at 1 kHz; average output 42 mW; Rapp OptoElectronic, Germany) was integrated with the imaging platform to deliver spatially confined injury at user-defined coordinates. Ablation was performed at 70% of maximum output power, with the beam guided along a short linear trajectory corresponding to the predicted plaque-ulceration axis. Laser settings and trace length were held constant across experiments to ensure comparable injury burden.

These parameters reproducibly generated loss of approximately 5–7 endothelial cells at the target site without disrupting the underlying PDMS–glass architecture. Chips exhibiting cell removal outside this range were excluded prior to perfusion. Immediately after ablation, channels were perfused with recalcified whole blood under the patient-specific flow conditions described above. Platelet recruitment, thrombus development, and embolic detachment were recorded in real time by confocal microscopy as detailed in the imaging section.

For thrombosis experiments, citrated whole blood was loaded into 5 mL Terumo syringes and perfused through the artery-on-chip devices using a Legato® 111 dual infuse/withdraw programmable touch-screen syringe pump (KD Scientific).

### 4.6 Pharmacological perturbations

Inhibitors were added to whole blood 10 min before perfusion. Concentrations: 5D2 and 6G1 (25 μg/mL, gifts from Dr. Timothy Springer); ARC1172 (1 μM, synthesized by Integrated DNA Technologies); caplacizumab (1 μM, Sigma); tirofiban (0.5 μM, Sigma); 152B6 (25 μg/mL, Emfret Analytics); LJ-P5 (25 μg/mL, Kerafast). Doses were chosen based on published studies (Table 2).

### 4.7 Image analysis and statistics

Calculation of platelet fluorescence intensity, coverage area, and embolisation were conducted using custom-made ImageJ programs. Fluorescence intensity over time was extracted from time-stack images of each artery-chip experiment after image processing to remove noise fix alignment issues. The intensity data was then saved as csv files for further analysis. Platelet area over time were obtained using the ImageJ plugin Trackmate with custom-made parameters. Changes in platelet area and displacement over time were used to determine embolisation events manually.

Each of these algorithms were compiled into batch processing packages to process experimental files efficiently. Graphed figures of fluorescence intensity, platelet area, and number of embolisations were made using the data collected and the PRISM software for clear visualisation and comparison of results. *p < 0.05, **p < 0.01, ***p < 0.001, ****p < 0.0001.

### 4.8 Molecular dynamics simulations

The VWF mechanomodule spanning the D′D3–A3 domains was modelled by superimposing the crystal structures of D′D3 (PDB: 6N29), A1 (PDB: 1SQ0), A2 (PDB: 3GXB), and A3 (PDB: 4DMU)(*82–85*) onto a ColabFold-predicted scaffold(*86*). Overlapping regions were removed, retaining crystallographic cores connected by ColabFold-derived linker segments. Disialylated core-1 O-glycans (residues 1248, 1255, 1256, 1263, 1468, 1477, 1486, 1487, and 1679) and monosialylated bi-antennary N-glycans (residues 857, 1147, 1231, 1515, and 1574)(*87*) (Fig. S9 A) were modelled using Glycan Reader & Modeler in CHARMM-GUI (*88*) and incorporated into the atomistic model. All simulations were performed using GROMACS 2021.2(*89*) with the CHARMM27 force field and TIP3P water. Systems were solvated, neutralized with Na⁺/Cl⁻ ions, energy-minimized by steepest descent, and equilibrated under NVT and NPT ensembles (100 ps each) at 300 K using a 2 fs time step. Production simulations (100 ns) employed particle-mesh Ewald electrostatics and optimized nonbonded cutoffs. Simulations exhibiting diverging backbone RMSD were restarted from the final frame until convergence to render the bird’s nest conformation of VWF (Fig. S9B and C). The equilibrated compact “bird’s-nest” conformation was placed in an elongated simulation box (15 × 15 × 150 nm) and subjected to flow molecular dynamics by pulling solvent toward the protein using an umbrella sampling bias. Cα atoms of the D′D3 domain were harmonically restrained (1000 kJ mol⁻¹ nm⁻²) to prevent translational drift, while only water oxygen atoms were pulled to minimize high-frequency noise. Flow was applied by pulling a 5-nm slab of water oxygen atoms along the Z-axis using a harmonic spring (1000 kJ mol⁻¹ nm⁻²) at a constant velocity of ∼0.01 nm ps⁻¹. Although exceeding physiological shear rates, this acceleration enables observation of unfolding within accessible simulation timescales(*90*). Simulations were performed in triplicate. Structural and dynamical analyses were performed using GROMACS tools and MDTraj(*91*) trajectories were visualized in PyMOL and QtGrace(*92, 93*). Flow MD trajectories (n = 3) were analyzed against a static reference containing ARC1172 (PDB: 3HXO)(*77*) and caplacizumab monomer (PDB: 7EOW)(*94*), due to unresolved structures of the dimeric caplacizumab in complex with VWFA1 (Fig. S9D) . The VWFA1 domain was aligned using heavy atoms of residues 1290–1430. Interatomic distances were calculated between heavy atoms of the static drugs and dynamic regions of interest, including the N-terminal AIM (NAIM; residues 1238–1271) and C-terminal AIM (CAIM; residues 1459–1493), together with proximal glycans within 5 Å. Steric clashes were defined as heavy atom separations <3.0 Å and used to assess whether drug engagement requires mechanomodule unfurling and AIM exposure.

### 4.9 Scanning electron microscopy (SEM)

SEM was performed using a benchtop Phenom XL scanning electron microscope (Thermofisher Scientific, USA), on dried and fixed non-gold coated samples. Images were taken at 600 – 11000x magnification under low vacuum conditions (60 Pa) using a full back scatter detector configured with a working distance of ∼3.2 mm at 5keV.

## Acknowledgment

We thank Peter Su, Johnny Liu, Richard Tan and Yingqi Zhang for helpful discussion on microfluidic imaging and thrombosis diagnosis. We thank Mike Wu, David Robinson, Gemma Figtree, Michael Kassiou, Ben Freedman, and Vivien Chen for clinical guidance and support for providing critical suggestions. This work was supported by the National Health and Medical Research Council (NHMRC) of Australia (APP2047049 and APP2047218 – L.A.J.; GNT2022247 – Y.C.Z.); NSW Cardiovascular Capacity Building Program (Early-Mid Career Researcher Grant – L.A.J.); MRFF Cardiovascular Health Mission Grants (MRF2023977 – L.A.J., MRF2016165 – L.A.J.); MRFF Early to Mid Career Researchers Grant (MRF2028865 and MRF2037779 – L.A.J.); Ramaciotti Foundations (2020HIG76 – L.A.J.); National Heart Foundation (106979 – L.A.J.; 106879 – Y.C.Z.), Office of Global and Research Engagement (USYD-UCL Ignition Grant – L.A.J.), and University of Sydney Drug Discovery Initiative (Cardiovascular Priority Driven Program – L.A.J., M.M.K. and F.P.). Lining Arnold Ju is a National Heart Foundation Future Leader Fellow Level 2 (105863) and a Snow Medical Research Foundation Fellow (2022SF176). All data that support the findings of this study are available on request from the corresponding author.

## Author contributions

L.A.J. and Y.C.Z. designed the research, analyzed data and wrote the manuscript. Y.C.Z designed the chip, cultured the endothelial cells and functionalized the chips, performed blood perfusion, conducted analysis and interpretation of data; Y.L. cultured the endothelial cells and functionalized the chips, conducted analysis and interpretation of data Z.W. designed and fabricated the chips, wrote the manuscript, and performed data analysis. N.L. performed the molecular dynamics simulation and molecular docking. A.N. performed immunostaining and data analysis; A.S. performed the CFD simulation and data analysis; Y.C. designed the anti-VWF analysis and wrote the manuscript. A.D. performed the immunostaining and conduct analysis, E.M.O. neutralized the collagen, M.M.K., K.B., X.X., J.Y., F.P., and T.A. provided clinical suggestions, co-wrote and revised the paper. T.A. provided the CTA images and contributed the preliminary data. L.A.J is the corresponding author.

