## Supplementary Information for "Patient-specific "Physical Twin" artery-on-chip platform reveals complex flow-dependent VWF mechanobiology and guides personalized antithrombotic therapy"

### Supplementary Figures

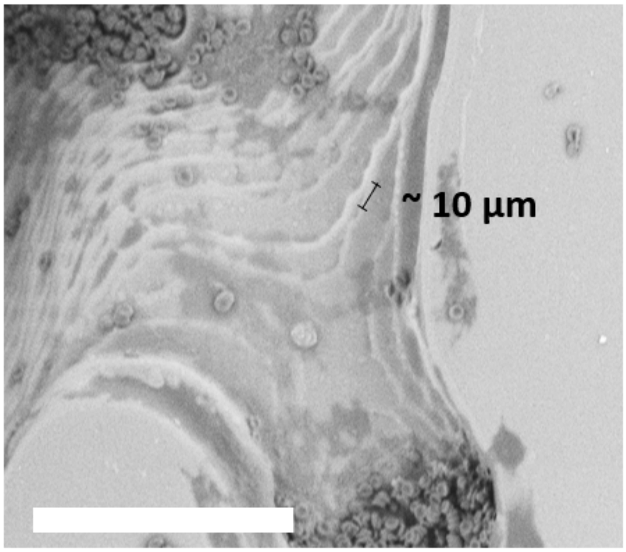


**Supplementary Figure S1. Scanning electron microscopy validation of printing resolution.** Scanning electron micrograph of a representative microchannel wall produced using the rapid glass-substrate digital light printing process, with a pixel resolution of approximately 10 µm. Scale bar, 80 µm.

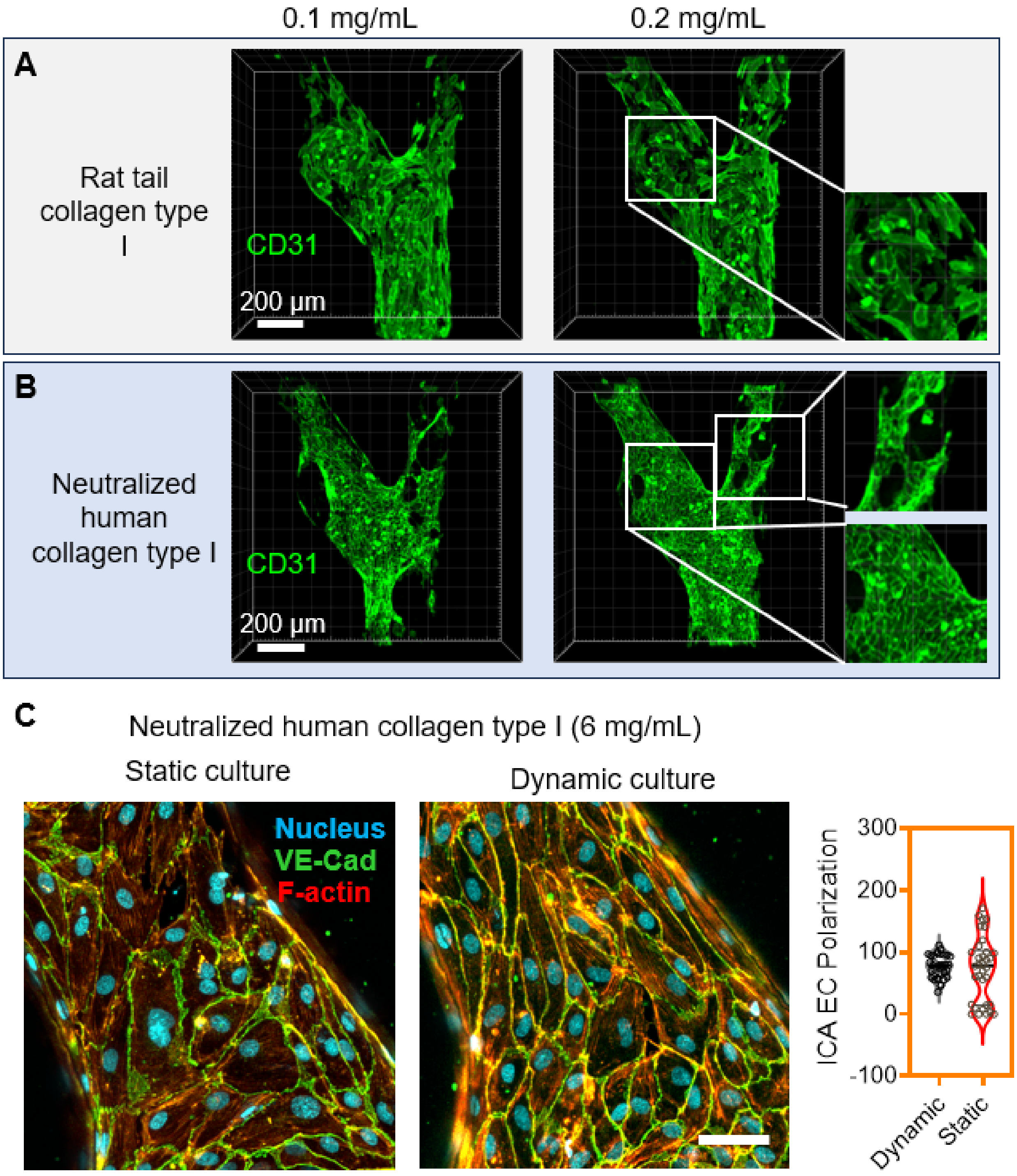


**Supplementary Figure S2. Neutralized human collagen type I supports endothelial junction formation in microfluidic carotid geometries.** Representative 3D confocal reconstructions of HCtAEC monolayers cultured within a patient-specific bifurcated artery-on-chip after coating with type I collagen at the indicated concentrations. Endothelium is visualized by CD31 immunostaining (green). **(A)** Rat-tail collagen type I coatings (0.1 and 0.2 mg/mL) supported cell attachment but produced heterogeneous coverage and discontinuous junctional organization, with enlarged views highlighting patchy CD31 signal and impaired monolayer maturation. **(B)** Neutralized human collagen type I coatings (0.1 and 0.2 mg/mL) promoted more uniform endothelialization and continuous CD31-positive cell–cell borders, consistent with improved barrier formation; inset magnifications show dense, junctional CD31 enrichment compared with rat-tail collagen. Scale bars, 200 µm. **(C)** Representative high-magnification immunofluorescence images of HCtAECs cultured on neutralized human collagen type I (6 mg/mL) under static or dynamic flow-conditioning conditions. Nuclei (cyan), VE-cadherin (green), F-actin (red). Dynamic culture promotes elongation and alignment of endothelial cells, consistent with a polarized arterial phenotype, whereas static culture yields a less ordered morphology. *Right*, quantification of ICA endothelial cell polarization. Scale bar = 30 µm. Panels A and B represent preliminary optimization experiments using sub-optimal coating concentrations (0.1 and 0.2 mg/mL).

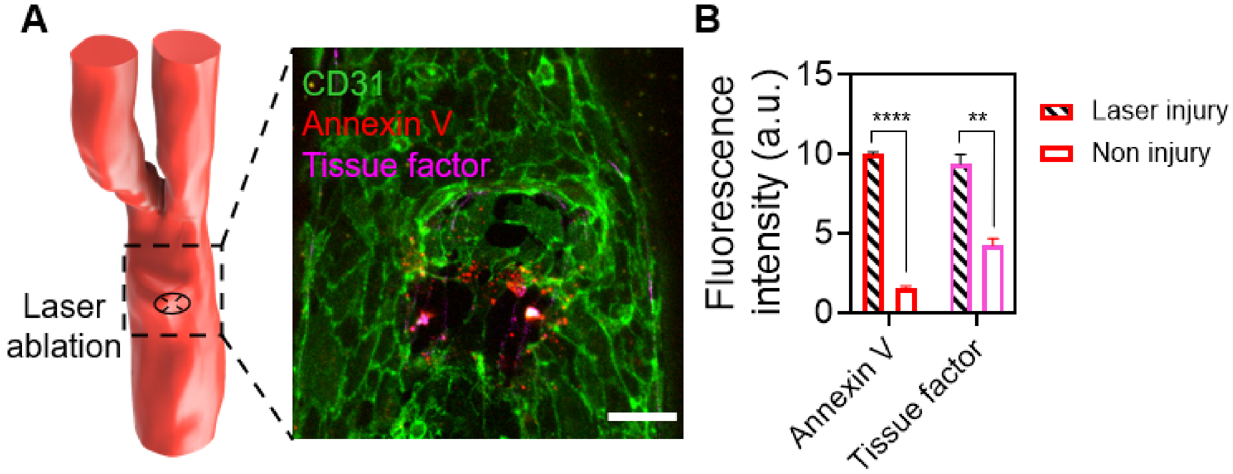


**Supplementary Figure S3. Injury-responsive phenotype of laser-ablated HCtAECs. (A)** Schematic and representative confocal image showing the laser injury site within the endothelialized channel. HCtAECs are labeled with CD31 (green), while Annexin V (red) and tissue factor (magenta) mark injury-associated pro-thrombotic responses. **(B)** Quantification of fluorescence intensity demonstrates significant enrichment of Annexin V (****$p < 0.0001$) and tissue factor (**$p < 0.01$) at laser-injured regions compared with non-injured controls, confirming that HCtAECs mount a localized pro-thrombotic injury response.

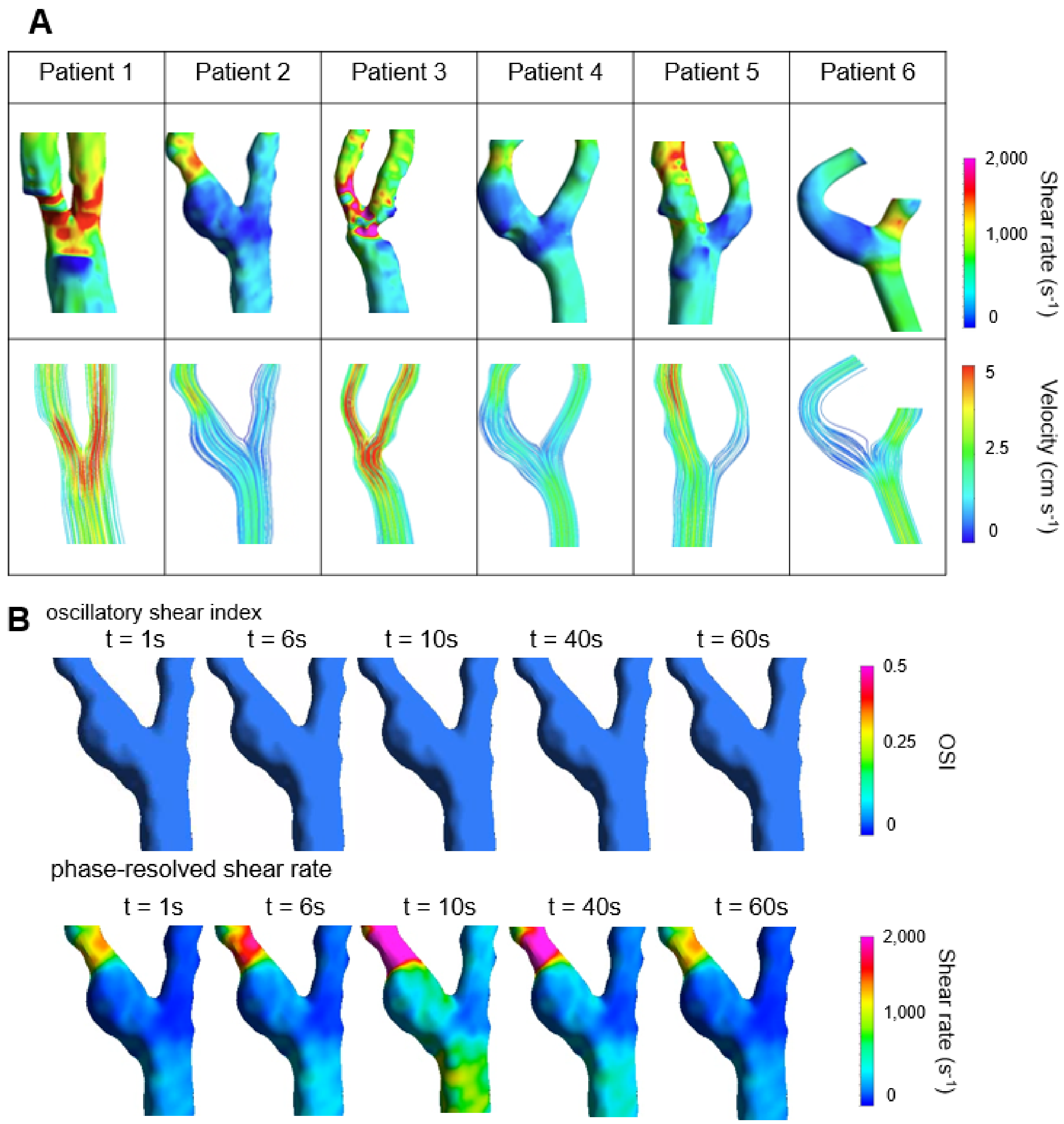


**Supplementary Figure S4. Computational fluid dynamics atlas of patient-specific carotid geometries.** **(A)** Comparative CFD analysis of six patient-derived carotid artery reconstructions used to generate the Physical Twin chips. **Top row:** surface maps of wall shear rate ($s^{-1}$) across each geometry, demonstrating marked inter-patient heterogeneity in both magnitude and spatial distribution of hemodynamic stress. **Bottom row:** velocity streamline fields (cm $s^{-1}$) highlighting geometry-dependent flow patterns. **(B)** Representative time-dependent CFD snapshots showing oscillatory shear index and phase-resolved wall shear-rate distributions under the measured pulsatile inlet condition.

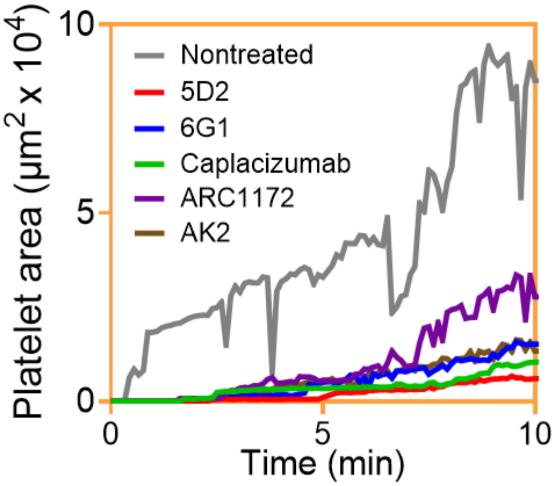


**Supplementary Figure S5. Thrombus area kinetics (high shear).** Untreated controls exhibit rapid exponential growth. All VWF–GPIbα–targeted agents markedly suppress area expansion, with 5D2 and caplacizumab producing the strongest inhibition.

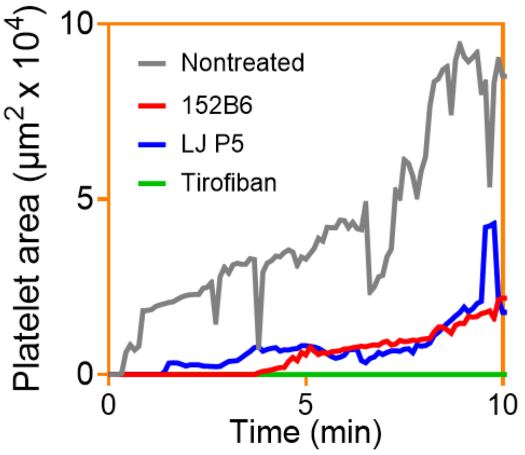


**Supplementary Figure S6. Integrin-pathway inhibition in the high-shear patient model.** Quantification of platelet-covered area over 10 min demonstrates that disruption of the fibrinogen-$\alpha_{IIb}\beta 3$ axis with tirofiban nearly abolishes thrombus growth, while selective interference with VWF-$\alpha_{IIb}\beta 3$ interactions (152B6, LJ-P5) produces only partial reduction

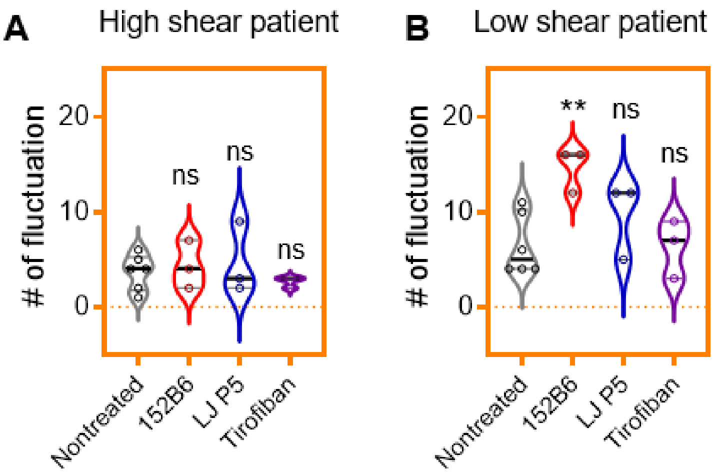


**Supplementary Figure S7. Quantification of platelet intensity fluctuations as a metric of thrombus instability. (A)** Number of platelet fluorescence intensity fluctuations in the high-shear patient model under the indicated inhibitor conditions. No significant differences were observed among treatments. **(B)** Number of platelet fluorescence intensity fluctuations in the low-shear patient model. Treatment with 152B6 significantly increased fluctuation frequency (**$p < 0.01$) relative to untreated controls, indicating enhanced thrombus instability, whereas LJ-P5 and tirofiban did not significantly alter this metric. Fluctuation events were defined from the platelet fluorescence time traces as each decrease of 1 a.u. in mean intensity, providing a quantitative readout of repeated thrombus destabilization and partial fragmentation.

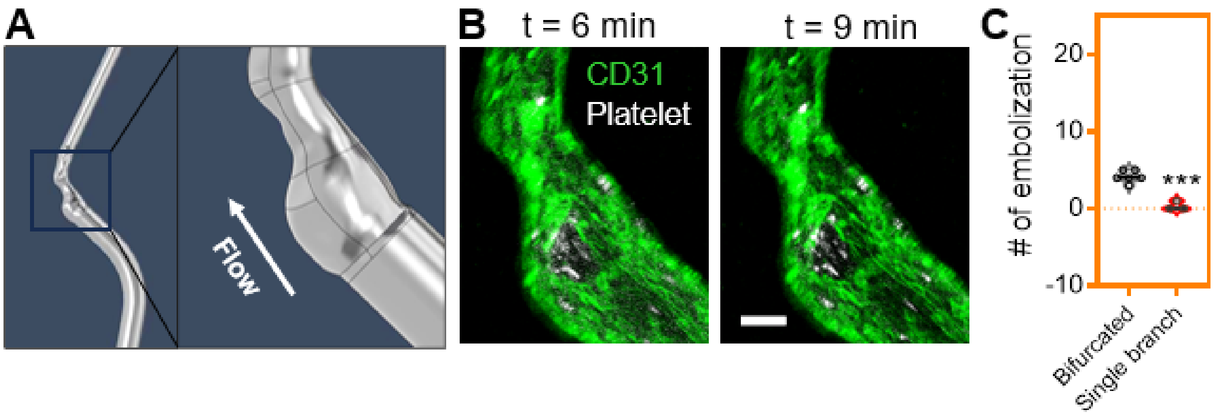


**Supplementary Figure S8. Single-branch model reveals the necessity of full bifurcation geometry for reproducing embolization (A)** Design of a simplified microfluidic construct representing only the internal carotid artery (ICA) branch from the low-shear patient, fabricated to preserve the local shear rate at the injury site while eliminating the bifurcation and upstream flow separation present in the full patient geometry. The zoomed inset illustrates the smooth, non-bifurcating lumen used for comparative testing. **(B)** Time-lapse confocal images of thrombosis after laser injury in the single-branch model show formation of a localized platelet aggregate (white) on intact endothelium (CD31, green) with markedly reduced fragmentation compared with the bifurcated counterpart. Scale bar, 80 μm. **(C)** Quantification of embolization events demonstrates a significant reduction in shedding in the single-branch configuration versus the full bifurcated geometry (***p < 0.001), despite matched local shear at the injury site.

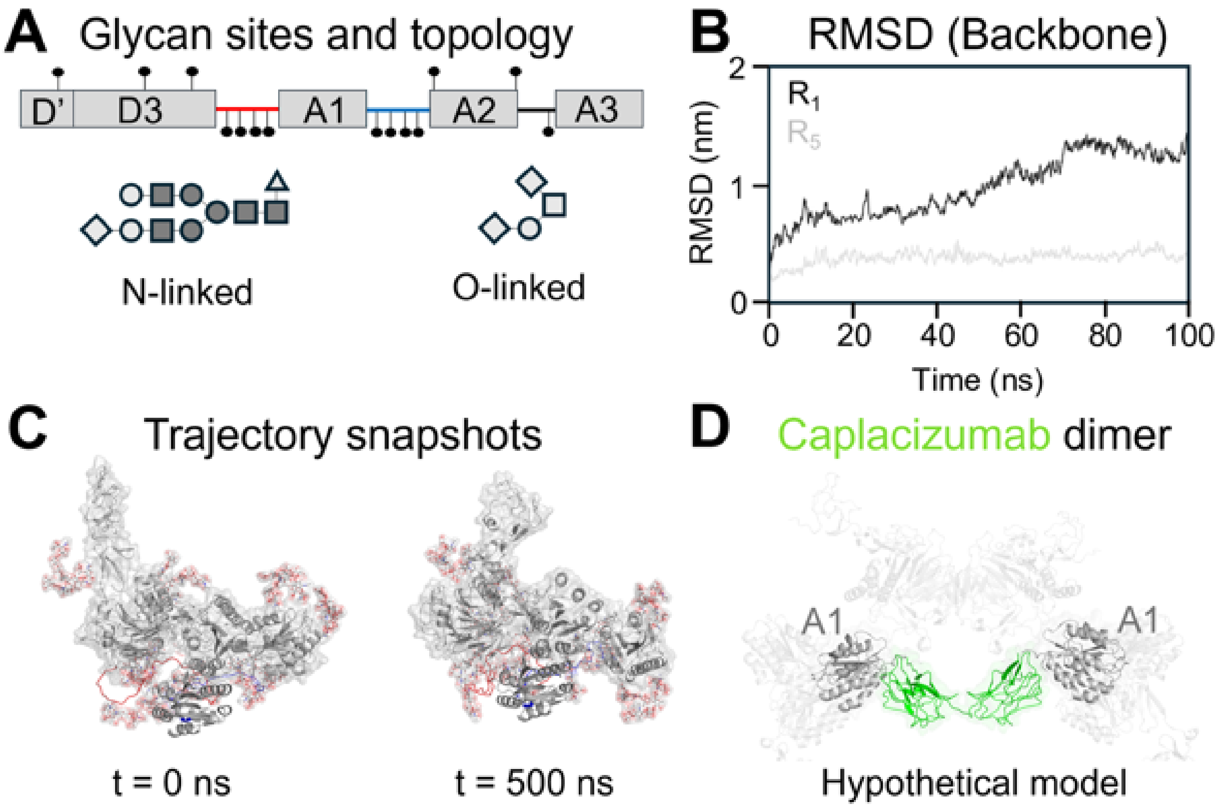


**Supplementary Figure S9. VWF Flow molecular dynamics settings. (A)** N- and O-linked glycosylation sites mapped across the VWF sequence and projected onto the D′ D3 – A1 – A2 – A3 mechanomodule, highlighting spatial distribution and topology. Glycan symbols follow: N-acetylglucosamine (dark grey square), N-acetylgalactosamine (grey square), galactose (grey circle), mannose (dark grey circle), fucose (grey triangle), and sialic acid (grey diamond). **(B)** Backbone RMSD of the VWF mechanomodule during equilibration, comparing early (0–100 ns;

R1) and late (400–500 ns; R5) time windows. **(C)** Representative snapshots of the pre-equilibrated (t = 0 ns) and equilibrated (t = 500 ns) structures, showing adoption of the compact “bird’s-nest” conformation. **(D)** Hypothetical model of Caplacizumab dimer.

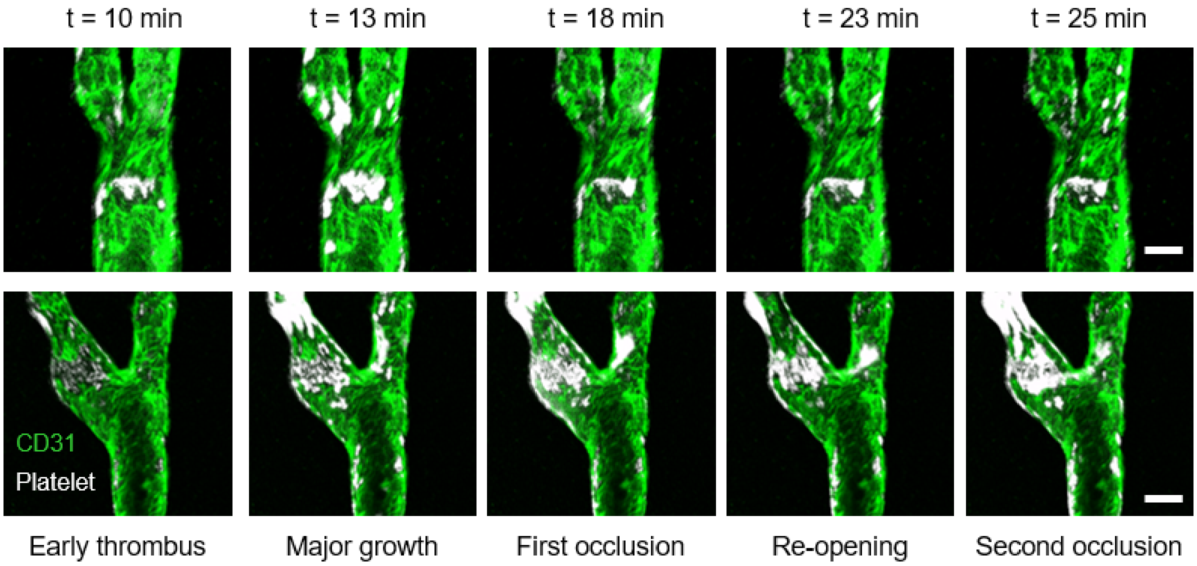


**Supplementary Figure S10.** Representative time-lapse confocal images from extended whole-blood perfusion experiments in the high-shear (top row) and low-shear (bottom row) patient-specific artery-on-chip models following laser-induced endothelial injury. In both geometries, early thrombi formed within the first 10 min and continued to enlarge during prolonged perfusion. By ~18 min, the developing aggregates reached a first occlusive state, followed by transient re-opening / reperfusion before progression to a second occlusion. Scale bars, 100 µm (top); 200 µm (bottom).
